# The Importance of Regulatory Network Structure for Complex Trait Heritability and Evolution

**DOI:** 10.1101/2024.02.27.582063

**Authors:** Katherine Stone, John Platig, John Quackenbush, Maud Fagny

## Abstract

Complex traits are determined by many loci—mostly regulatory elements—that, through combinatorial interactions, can affect multiple traits. Such high levels of epistasis and pleiotropy have been proposed in the omnigenic model and may explain why such a large part of complex trait heritability is usually missed by genome-wide association studies while raising questions about the possibility for such traits to evolve in response to environmental constraints. To explore the molecular bases of complex traits and understand how they can adapt, we systematically analyzed the distribution of SNP heritability for ten traits across 29 tissue-specific Expression Quantitative Trait Locus (eQTL) networks. We find that heritability is clustered in a small number of tissue-specific, functionally relevant SNP-gene modules and that the greatest heritability occurs in local “hubs” that are both the cornerstone of the network’s modules and tissue-specific regulatory elements. The network structure could thus both amplify the genotype-phenotype connection and buffer the deleterious effect of the genetic variations on other traits. We confirm that this structure has allowed complex traits to evolve in response to environmental constraints, with the local “hubs” being the preferential targets of past and ongoing directional selection. Together, these results provide a conceptual framework for understanding complex trait architecture and evolution.

## Introduction

In many organisms, including yeasts, insects, worms, plants, and mammals, adaptation often leverages polygenic traits to respond to new environmental challenges(1–3). Such “complex” traits have an interconnected genetic architecture involving from a small number to potentially thousands of partly independent loci(4–7). Up to 90% of the SNPs associated with such traits are localized outside of coding regions, according to genome-wide association studies (GWAS) (8, 9). They are thus likely to affect traits by modifying gene expression—often by altering the sequence of *cis*-regulatory elements. This is consistent with the observation that GWAS-significant regions co-localize with SNPs that are linked to the expression of nearby genes (expression quantitative trait loci, *cis-*eQTLs) (10, 11). Partitioning heritability for these traits across various annotations while correcting for linkage disequilibrium has confirmed that *cis*-eQTLs as a group explain more of complex trait heritability than would be expected by chance (12–14) but still fail to explain a majority of trait heritability (15). Others have shown that including *trans*-eQTLs may capture additional heritability and explain important pathological mechanisms (16), but this is rarely done because *trans*-eQTL studies are generally underpowered.

Advances in systems biology and functional genomics have highlighted the fact that at the molecular level, complex traits are the results of many regulatory interactions between actors of different biological processes within a complex cellular gene regulatory network (17, 18). This high level of pleiotropy coupled with significant epistatic interactions may explain the missing heritability observed in quantitative genetics studies and has led to the development of the omnigenic model (19, 20). In this model, trait association signals are spread across most of the genome in a way that includes many genes lacking an obvious connection to a particular trait. The model posits that most heritability can be explained by effects on genes outside of “core pathways,” often acting in *trans*, but acting in ways that alter the functioning of those pathways and thus account for most of a given trait’s heritability. The amount of pleiotropy implied by this model may limit, at first glance, the possibility for a given complex trait to adapt in response to an environmental change without interfering with other unrelated traits. However, the evidence for polygenic adaption processes highlights the need to dig deeper into the molecular bases of complex traits to understand their architecture and how they adapt (21).

Given the number of loci and the number of potential interactions involved, gene regulatory networks provide an efficient tool for understanding the biological processes behind trait heritability (22). We previously proposed a way to integrate both *cis*- and *trans*-eQTL results using a bipartite eQTL network representation (23, 24), that relies on summary statistics from eQTL studies and is relatively insensitive to a high false discovery rate, allowing one to partially compensate for the lack of power of *trans*-eQTL studies; an update to this method allowed us to weigh SNP-gene regulatory relationships by eQTL effect sizes (25). These analyses have shown that highly structured eQTL networks can reliably identify, in a tissue-specific manner, the biological functions disrupted by traits-associated SNPs (23, 26), and further defined network topological features that are useful in explaining in part the link between genotype and phenotype in complex traits.

In this study, we build on eQTL networks to investigate how complex trait heritability is spread within and among the tissue-specific eQTL networks to better understand both the genetic architecture of complex traits and how they may evolve under directional selection. We used GWAS summary statistics from ten complex traits and diseases and built 29 tissue-specific *cis*- and *trans*-eQTL networks using the GTEx dataset. We then partitioned heritability across various features, including network node topological summary statistics, to identify key determinants of trait heritability. We investigated whether heritability for each trait is clustered in particular subparts (regulatory modules, also sometimes referred to as communities) of the eQTL networks, and identified network communities regulating biological functions that explain most of the heritability of each trait. Finally, in order to assess how complex traits may evolve, we looked for signatures of past polygenic adaptive processes in key regulatory loci considered in relationship with their role in the eQTL network. We found that heritability is not scattered uniformly across the genome but rather “clustered” in eQTL modules that represent trait-relevant functions and that loci playing a key role in regulating tissue-specific biological processes were not only more likely to explain trait heritability but also to have been preferential targets of past polygenic adaptation events.

## Material and Methods

### GTEx data set

We used genotyping and gene expression level data from the NHGRI GenotypeTissue Expression (GTEx) project version 8.0 (27). For appropriate statistical power in downstream analyses and network stability, we filtered out tissues for which the number of individuals with both genotyping and RNA-Seq data available was less than 200; sex-specific tissues were not included. This left 29 tissues for analyses, as can be found in Supplementary Tab. S1. Genotyping data were downloaded from the database of Genotypes and Phenotypes (dbGaP): phs000424.v8.p2. Genotyping data were preprocessed on the Bridges system at the Pittsburgh Supercomputing Center (PSC) and the Cannon cluster supported by the Faculty of Arts and Sciences Division of Science, Research Computing Group at Harvard University (see (25)).

The sequencing data were processed in plink 1.90 to retain only SNPs, and we removed variants with genotype missingness greater than 10% or minor allele frequency less than 0.1 (28). SNP imputation was then performed using Eagle2 (29).

Fully processed, filtered, and normalized RNA-Seq data were obtained from the GTEx Portal (www.gtexportal.org). The GENCODE 26 model was used to collapse transcripts and quantify expression using RNA-SeQC (https://www.gencodegenes.org/human/release_26.html#).

### Bipartite eQTL networks inference

Expression quantitative trait loci (eQTLs) were obtained from (25). Rapidly, with the R MatrixEQTL package (30), the association between SNP genotypes and gene expression was modeled using linear regression (Eq. 1) that included potential confounding factors as covariates (31): the two first principal components for population structure, sex, age and RIN that measures RNA quality. If **G** is an *r×m* matrix of gene expression and **S** is an *r×n* matrix of SNP genotypes, each with *r* rows representing observations and columns representing *n* SNPs and *m* genes, respectively, **X** is a covariate matrix, the eQTL of a particular SNP *i* on a locus’s gene expression *j* is then :

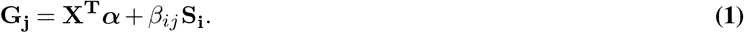

Associations were evaluated for SNPs in both *cis* – SNPs within 1MB of a gene’s transcription start site – and *trans*. The eQTL associations between all pairs of SNPs and genes were then represented as a sparse, weighted bipartite network (Fig. 1). Each SNP and gene was considered a node in the network. Using a fixed cutoff *q* = 0.2 on the false discovery rate (FDR) of the eQTL regression (23, 24), the edge weight *a*_*i,j*_ between SNP *i* and gene *j* were defined by the function 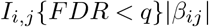. Thus, when the estimated FDR of the eQTL regression was below the threshold of 0.2, then 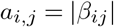, indicating there was an edge connecting the nodes, and *a*_*i,j*_ = 0 otherwise.

**Fig. 1.**
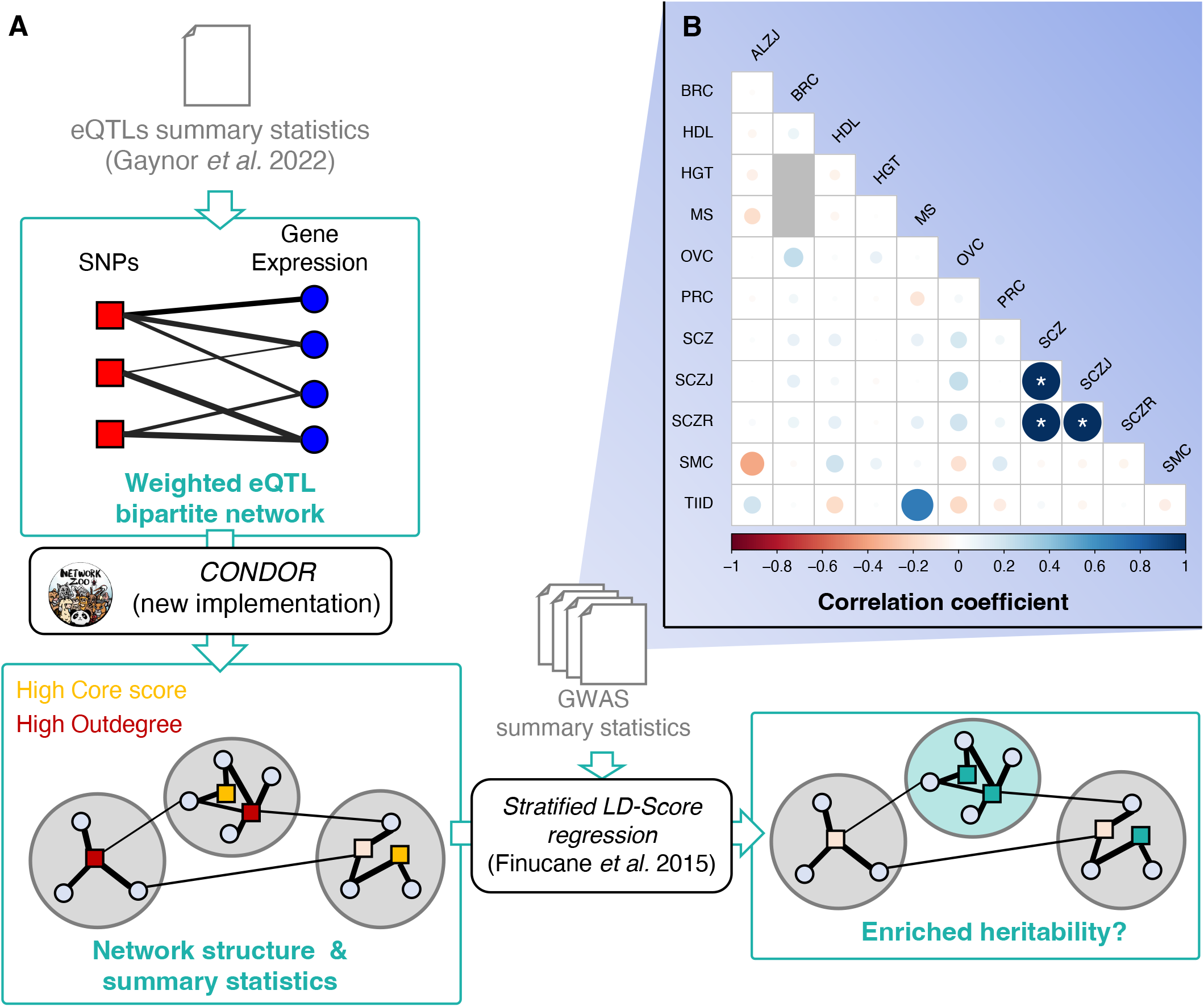
Pipeline and Traits correlation. **A**. Pipeline of data analyses. Input data are in grey. GTEx eQTL summary statistics were obtained from a previous study (25) and are described in Methods and Supplementary Tab. S1. GWAS summary statistics were downloaded from https://console.cloud.google.com/storage/browser/broad-alkesgroup-public-requester-pays and are described in Supplementary Tab. S2. eQTL network summary statistics are presented in Supplementary Fig. S1. **B**. Pairwise genetic correlations between traits based on GWAS summary statistics.

### Network summary statistics

The module structure of each tissue-specific network was determined using the bipartite modularity maximization approach (24), allowing for a balance between computation time and memory usage (Fig. 1). This new implementation (CONDOR:condorSplitMatrixModularity) has been published in the R netzooR package fork of https://github.com/maudf/.

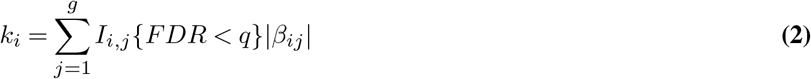

The SNP core score was defined as the SNP’s contribution to the modularity of its module, it measures the centrality of the SNP in the module (24). If *m* = *s×g* is the total number of possible edges in a network made of *s* SNPs and *g* genes, 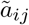 is the observed edge value between SNP *i* and gene *j*. Here, *d*_*j*_ is the gene *i* indegree defined as 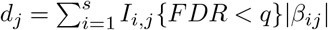, then, for SNP *i* in module *h*, its core score, *Q*_*ih*_, is defined by Eq. 3:

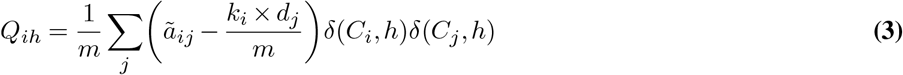

### GWAS data

GWAS data were obtained from the Alkes group (see Supplementary Tab. S2). We considered a total of ten traits and diseases presenting varying levels of estimated genetic heritability. For each trait or disease, we obtained summary statistics including SNP chromosome, position, allele 1 and 2, *χ*^2^, and *Z*-score. Here, the relationship between an individual’s *i* genotype at SNP *s* and its phenotype *Y*, accounting for covariates *M*, is usually tested using the following regression: *Y*_*i,s*_ = *β*_1,*s*_*X*_*i,s*_ + *β*_2_*M*_*i*_ + *ϵ*_*i,s*_, were *ϵ*_*i,s*_ is the error term. Then 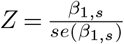 and *χ*^2^ = *qnorm*((*P/*2)^2^) where *P* = 2 *×pnorm*(−|*Z*|). We used *Z*^2^ as a proxy for normalized heritability explained by each SNP. High heritability SNPs were defined as those with a *Z*^2^ in the 95th percentile of the distribution.

### GWAS Outdegree and Core Score enrichment among high heritability SNPs

We compared the distribution of SNP outdegrees and core scores between the high heritability SNPs and the rest of the SNPs in the network using a likelihood-ratio test (LRT), correcting for linkage disequilibrium. To control for LD between SNPs, we generated lists of SNPs falling into the same LD block, using the plink1.9 –blocks option, a 5-Mb maximum block size, and an *r*^2^ of 0.8. In each module, for each LD block, we extracted the median of either outdegrees (*k*_*i*_) or core scores (*Q*_*ih*_) for high and non-high heritability SNPs separately and used these values as input in the linear regressions.

The LRT we used assesses whether a linear model that includes GWAS status (Eq. 5) fits the observed data better than a linear model that does not include this variable (Eq. 4). As the distribution of SNP *Q*_*ih*_ is not uniform across modules, we added module identity as a covariate in the linear regression when computing LTR for core scores. In Eqs. 4 and 5, *Score*_*i*_ is the score of SNP *i* (either outdegree or core score), *I*(*GWAS* = 1) is an indicator function equal to 1 if the SNP is a high heritability SNP and equal to 0 otherwise, and *I*(*C*_*k*_ = 1) is an indicator function equal to 1 if the SNP belongs to module *k* and equal to 0 otherwise:

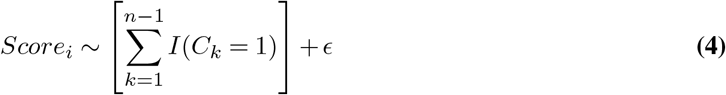

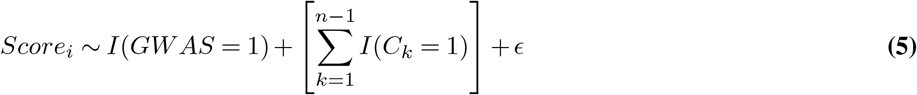

### Genetic correlation between traits and heritability enrichment analyses among local and global hubs

We computed partitioned heritability with the 97 annotation baseline-LD model from (32, 33) for each of the ten traits and diseases selected above (Fig. 1). This allowed us to estimate the enrichment and standardized effect size of the baseline annotations and three additional parameters on the heritability: belonging to the network (being an eQTL), having a high outdegree, and having a high core score (32, 33). We considered a binary annotation to ensure sufficiently stable estimates. For outdegrees and core scores, annotations were set equal to 1 if *k*_*i*_ and *Q*_*ih*_ were in the top quartile of the distribution and 0 otherwise.

Given that approximately 85% of the GTEx study population consists of individuals of European descent, we used LD scores computed from the 1000 Genomes Project data from individuals with European ancestry using the GRCh38 genome version and regression weights that exclude the HLA region; *p*-values were computed using a block-jacknife.

Meta-analyses were then performed across uncorrelated traits using the meta.summaries function from the R rmeta package v.3.0, with random-effect weights. To identify uncorrelated traits and diseases, pairwise genetic correlations between the ten traits and diseases were first computed using stratified LD score regression (S-LDSC). Traits and diseases with a pairwise genetic correlation of less than 0.3 were considered uncorrelated.

### High heritability SNP enrichment among modules

High heritability SNP enrichment among modules was performed using a *χ*^2^-test on data corrected for linkage disequilibrium. Using the same LD-blocks as previously described, we considered that an LD block was a high heritability block if at least one SNP in this LD block had a *Z*^2^ in the top 95th percentile. For each module, we counted the number of high and non-high heritability LD blocks in and outside of the modules and used these data to perform a *χ*^2^-test.

### Gene Ontology enrichment analyses and identification of tissue-specific modules

We performed Gene Ontology enrichment analyses using the Bioconductor R topGO package v.2.44 (34), using the elim method; this method is more efficient than Fisher’s Exact test (35). The GO categories are tested sequentially, following the GO tree structure from bottom to top: if one GO category is found to be significant, the genes involved are removed from the parent nodes before they are tested. The tests are thus not independent, and no multiple testing correction can be applied. Following the guidelines in the topGO users’ manual, we filtered uncorrected p-values using a stringent threshold of 0.01. We also filtered out GO categories that did not include at least three genes in the gene set of interest. The gene ontology database used in this analysis was the one from the R bioconductor org.Hs.eg.db package v.3.13.0. For each test, the background gene set contains all the genes of the network and the gene set of interest contains all the genes from the module of interest.

We identified common and tissue-specific modules in the eQTL networks based on pairwise comparisons of GO Term Assignments. For each module of a first network, the GO ID enriched in the module was compared to the GO ID enriched in each of the modules in a second network using Jaccard Index. The best matching module was determined based on the highest Jaccard Index, and if the best Jaccard Index was ≥ 0.3, the modules were considered as similar and otherwise different. Then, for each module in each tissue-specific network, we counted the number of similar modules in the other 28 networks. If this number of similar functional modules was ≤ 2, we considered it a tissue-specific module.

### Selection scores enrichment analyses

Conserved SNPs were determined by intersecting the annotations of the SNPs present in our eQTL networks with conserved regions determined using GERP scores downloaded from Ensembl 111 release (https://ftp.ensembl.org/pub/release-111/bed/ensembl-compara/91_mammals. gerp_constrained_element/gerp_constrained_elements.homo_sapiens.bb (36)). Enrichment in conserved SNPs among local and/or global hubs were computed using the following logistic regression model, where *I*(*GERP* = 1) is an indicator function equal to 1 if the SNP falls in a conserved region and equal to 0 otherwise, *I*(*Cat* = 1) is an indicator function equal to 1 if the SNP belongs to a given category (local or global hubs) and equal to 0 otherwise, and *I*(*C*_*k*_ = 1) is an indicator function equal to 1 if the SNP belongs to module *k* and equal to 0 otherwise:

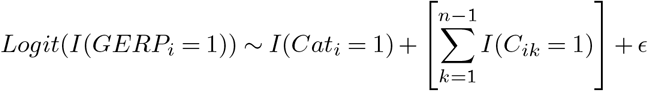

*iHS*, that measures the differences in haplotype lengths between haplotypes carrying the ancestral and the derived SNPs, detect signatures of recent selective sweeps (37). We downloaded *iHS* standardized scores (38) computed for 26 populations for 1000 genomes project Phase 3 dataset from https://zenodo.org/records/7842512. We extracted scores computed on the CEU population (Utah residents (CEPH) with Northern and Western European ancestry) samples as being the closest population from the European descent samples that represent 85% of the GTEx dataset v8 release.

*F*_*ST*_ between African, Eurasian and European samples from the 1000 genomes data were computed using plink2.0 (28) on genotypic data for samples from the EUR, AFR and EAS populations from the 1000 genomes Phase 3 dataset. Genotyping data were downloaded from ftp://ftp.1000genomes.ebi.ac.uk/vol1/ftp/data_collections/1000G_2504_high_coverage/working/20201028_3202_raw_GT_with_annot/.

Effect sizes of being a local or global hub on *iHS* or *F*_*ST*_ were computed using the following linear regression model, where *Score*_*i*_ is the value of the statistics for SNP *i, I*(*Cat* = 1) is an indicator function equal to 1 if the SNP belongs to a given category (local or global hubs) and equal to 0 otherwise, and *I*(*C*_*k*_ = 1) is an indicator function equal to 1 if the SNP *i* belongs to module *k* and equal to 0 otherwise:

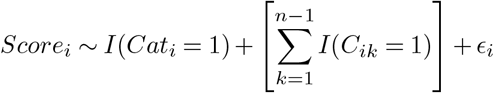

The community ID terms were only added when local hubs were one of the two categories compared, as the q-score determining local hubs depends on community ID.

## Data and code availability

All the code used to analyze the data is available at https://github.com/maudf/heritability. The new CONDOR:condorSplitMatrixModularity is available at https://github.com/maudf/netZooR The data used for the analyses described in this manuscript were obtained from: the GTEx Portal on 12/17/19 and dbGaP accession number phs000424.v8 on 12/17/19 for RNA-seq and Genotyping data, and from https://console.cloud.google.com/storage/browser/broad-alkesgroup-public-requester-pays for GWAS data.

## Results

### SNPs associated at the genome-wide level with traits or diseases are clustered in a few modules

We wanted to understand how heritability is distributed across the eQTL network. We had previously reported that SNPs significantly associated with chronic obstructive pulmonary disease, cancers, and other traits in GWAS are concentrated in a small number of modules (24) and wanted to verify that this result can be extended beyond genome-wide significantly associated SNPs to SNPs explaining a larger part of genetic heritability. We thus used RNA-seq and genotyping data from GTEx to perform eQTL analyses and built 29 tissue-specific weighted bipartite eQTL networks. We identified modules containing highly connected SNPs and genes within each network based on CONDOR’s bipartite modularity maximization (24) (see Methods, Supplementary Text and Supplementary Fig. S1 and Tab. S1).

We considered a total of ten traits and diseases chosen for their medical or evolutionary relevance in populations of European descent. Breast Cancer (BRC), Ovarian Cancer (OVC) and Prostate Cancer (PRC), Alzheimer’s Disease (ALZ), Multiple Sclerosis (MS), Schizophrenia (SCZ), high HDL levels (HDL), and Type 2 diabetes (TIID; diabetes) are all increasing in populations of European descent and are known to exhibit partial genetic heritability. Smoking Cessation (SMC) is a measure of smoking dependency that is partly explained by genetic factors and is linked to a host of pulmonary and other diseases. Schizophrenia (SCZ) is a highly heritable but poorly understood polygenic disease. Finally, Height (HGT) is the canonical example of a polygenic trait, and some consider it a natural selection target in populations of European descent. These traits and diseases also represent a wide range of global genetic heritability, as reported in the literature, from 25% for type 2 diabetes, to up to 80-90% for schizophrenia and Alzheimer’s. These traits and diseases also represent a wide range of global genetic heritability, as reported in the literature, from 25% for type 2 diabetes, to up to 80-90% for schizophrenia and Alzheimer’s. We conditioned on European descent because of the number of GWAS data available and the demographics of GTEx. A complete description of these traits and diseases and their GWAS summary statistics is reported in the Supplementary Tab. S2. For each trait or disease, we obtained summary statistics (see Methods).

We used the summary statistic *Z*^2^ as a proxy for the per-SNP heritability, and SNPs with a GWAS *Z*^2^ in the top 5% were named “high heritability SNPs.” We found that the high heritability SNPs cluster in a small number of network modules (Supplementary Tab. S3). As an example, the breast cancer-associated SNPs appear in only 29 (12.6%) of the 230 network modules in the SKN (Skin - Not sun-exposed–Suprapubic) eQTL network (Fig. 2A) representing a substantial concentration of heritability. We found similar results for other diseases and other tissues, with some combinations exhibiting even greater clustering of heritability in a limited number of network modules (Fig. 2B).

**Fig. 2.**
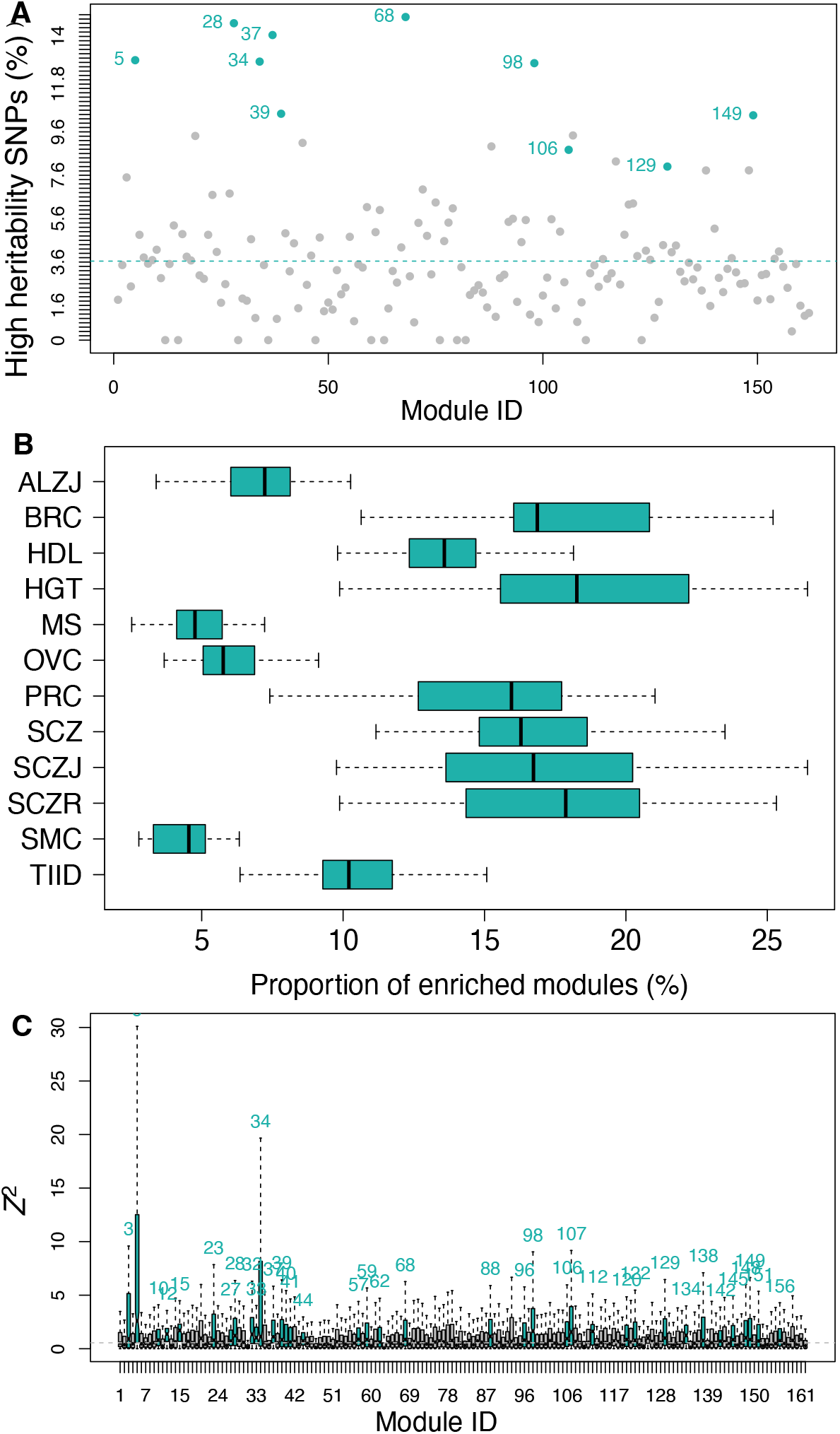
High heritability SNPs are clustered in a few trait-specific modules. **A**. Proportion of high heritability SNPs for Breast Cancer in each module of the Skin - Not sun-exposed (Lower Leg) eQTL network. Modules significantly enriched for high heritability SNPs are highlighted in blue (Benjamini-Hochberg corrected *p* ≤ 0.01 using a *χ*^2^-test). The dotted blue line represents the expected proportion of high heritability SNPs by chance in each module. **B**. Distribution of the proportion of modules enriched for high heritability SNPs for each trait or disease among all tissue-specific eQTL networks. **C**. Distribution of *Z*^2^ values in each module of brain – nucleus accumbens (basal ganglia) for Alzheimer’s disease (Kruskal-Wallis *p <* 2 × 10^−16^). Modules with a distribution skewed towards high values are represented in blue (modules with a Benjamini-Hochberg corrected *p* ≤ 0.01 using one-sided Mann-Whitney U tests for each module *vs*. the rest of the network were considered as significantly enriched in high *Z*^2^). The dotted blue line represents the expected *Z*^2^ median across the whole network.

Because the highest heritability SNPs often only explain a small proportion of the heritability of associated traits (particularly for highly polygenic traits), we examined the distribution of per-SNP heritability across the different eQTL network modules. We found that heritability was not distributed evenly across modules (Kruskal-Wallis test *p*=0 for all pairs of tissue-specific networks and traits tested). As an example, consider *Z*^2^ calculated for Alzheimer’s disease in Brain - Nucleus Accumbens (basal ganglia) networks, which is plotted in Fig. 2C; all the results are in Supplementary Tab. S3). Comparing the distribution of *Z*^2^ for each module with the rest of the network, we found that less than a third of the modules contain high heritability SNPs (54[5-84] modules representing about 23.7%[5.6-33.1] of the total number of modules depending on tissue-specific network/trait pair, with Benjamini-Hochberg-corrected Mann-Whitney U tests *p* ≤ 0.01 (see Supplementary Tab. S4).

### Heritability is enriched in trait-specific, functionally relevant modules

We investigated whether the heritability for different, uncorrelated traits was clustered in the same or different modules in the tissue-specific eQTL networks. Depending on the tissue-specific network, about 40% [28-53] of the modules were not enriched for high heritability SNPs associated with any traits, 29% [24-35] were enriched for high heritability SNPs from only one trait and can be considered trait-specific, and 3%[1-8] were enriched for high heritability SNPs from at least five of the ten traits and can be considered as shared across (many) traits (Fig. 3A and Supplementary Fig. S2).

**Fig. 3.**
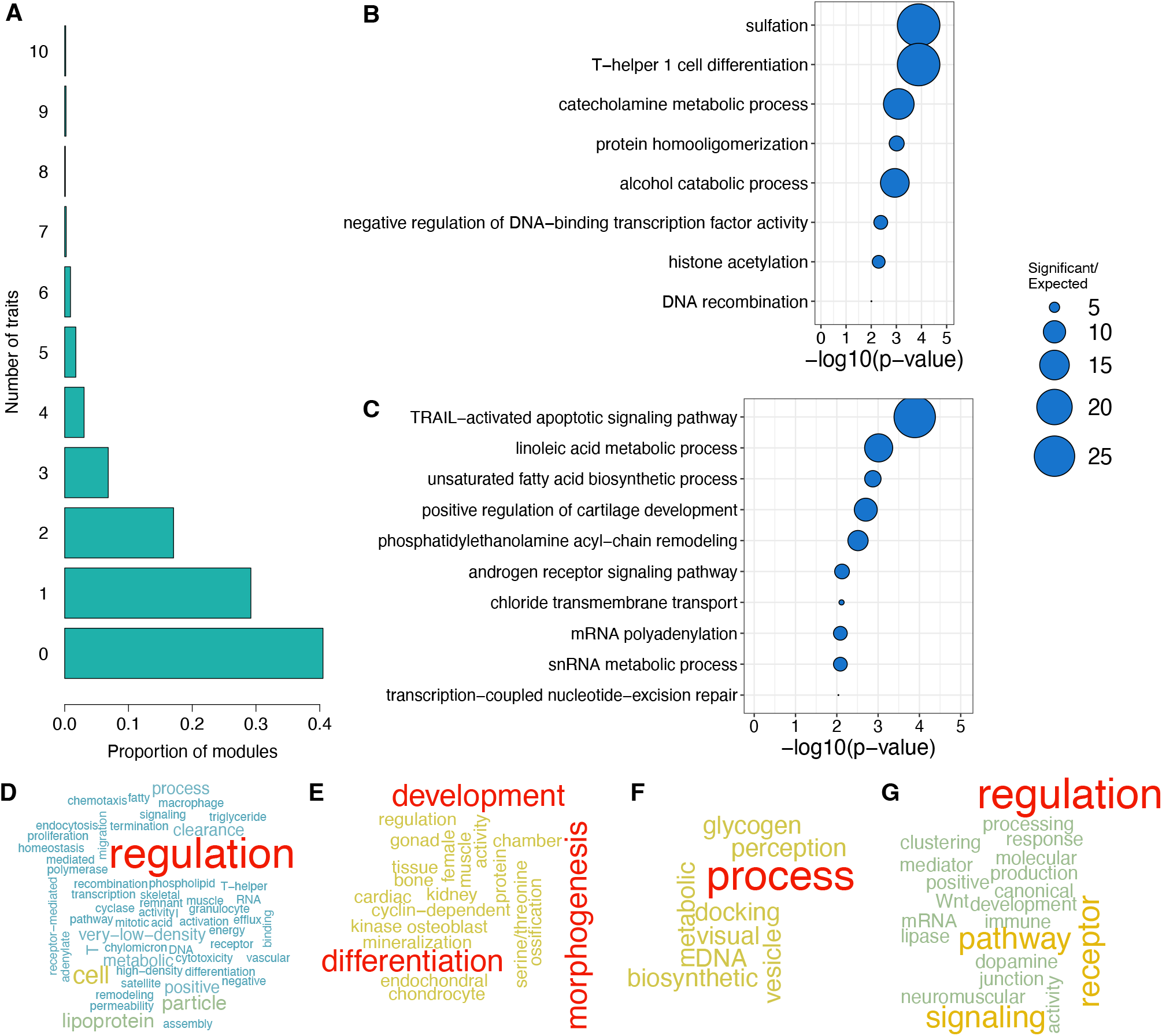
Heritability is clustered in trait-specific, biologically significant modules. **A**. Average proportion of modules enriched in high heritability SNPs for one to ten traits or diseases across all tissue-specific modules. To remove artifacts due to the high genetic correlation between the three schizophrenia studies, only one schizophrenia study was taken into account in the analysis. The 0 category represents the modules that are not enriched in high heritability SNPs for any trait or disease. For a split by tissue-specific network, see Supplementary Fig. S2. **B-G**. Gene Ontology enrichment analyses results on gene content for a few tissue-specific modules enriched for high heritability SNPs for one or two traits and diseases. **B-C**. Bubble plots for top ten terms or terms with p-value < 0.01 for **B**. Brain (nucleus accumbens - basal ganglia) module 5, enriched for high heritability SNPs for Alzheimer’s disease and multiple sclerosis. **C**. Colon - transverse module 149, enriched for high heritability SNPs for prostate cancer. **D-G**. Word clouds representing word frequencies in gene ontology terms enriched in **D**. Adipose - visceral omentum, module 142, enriched for high heritability SNPs for HDL; **E**. Muscle - skeletal, module 90, enriched for high heritability SNPs for height; **F**. Whole blood, module 17, enriched for high heritability SNPs for type 2 diabetes; **G**. Brain (nucleus accumbens - basal ganglia) module 100, enriched for high heritability SNPs for schizophrenia.

We performed a pairwise comparison of traits in each of the tissue-specific networks between modules enriched for high heritability SNPs. Using 10,000 resamplings, we found that top-heritability SNPs for two independent traits did not cluster in the same modules more than expected by chance in most cases. Indeed, among the 1305 pairwise comparisons possible 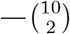 comparisons *×*29 tissues — only 28 show significant enrichment in overlap compared to what would expected by chance (Benjamini-Hocheberg-corrected *p* ≤ 0.01). In half of these cases, the excess of overlap was observed between high HDL (high-density lipoprotein), height, and/or and Type 2 Diabetes in various tissue-specific networks (see Supplementary Tab. S5). We also investigated the biological functions represented by genes with the modules enriched for high heritability SNPs. We performed a GO term enrichment analysis for each module using Bioconductor R topGO package (see Methods); the results are presented in Supplementary Tab. S6. This allowed us to identify modules that were tissue-specific (see Methods). We then focused on trait- and tissue-specific modules, defined as those enriched for high heritability SNPs from one or two traits and enriched for tissue-specific GO Terms (Supplementary Tab. S7)— in order to investigate the specificities of each trait and/or disease heritability and explicit the underlying molecular mechanisms. We found that these modules were enriched in genes involved in biologically and trait-relevant functions.

For example, heritability for Alzheimer’s disease and multiple sclerosis were both clustered in module 5 of the brain – nucleus accumbens (basal ganglia) network; module 5 is enriched in genes involved in catecholamine metabolic process (Fig. 3B), a class of molecules whose concentrations are altered in symptomatic Alzheimer’s and multiple sclerosis diseases, both of which are amyloid plaque diseases (39, 40). Heritability for schizophrenia was most strongly clustered in module 100 of the same tissue, a molecule that is enriched for dopamine receptor signaling pathway genes (Fig. 3G), and this pathway is known to be functionally disrupted in the brain striatum in schizophrenia patients (41).

Unsurprisingly, heritability for cancers, including breast cancer, ovarian cancer, and prostate cancer, were enriched in epithelial tissues (skin–not sun-exposed exposed, skin–sun-exposed, colon–transverse) in network modules consisting of genes enriched for immune response (prostate cancer and module 76 of skin, not sun-exposed), response to cellular hypoxia (breast cancer, prostate cancer and module 157 of skin–sun-exposed), DNA break repair, cell cycle, apoptosis, and epithelium differentiation and growth (breast cancer and module 149 of skin–sun-exposed, and module 191 of skin (not sun-exposed), ovarian cancer and module 199 and skin, prostate cancer and modules 78, 97 of skin and module 149 of skin–sun-exposed). Particularly interesting, prostate cancer heritability is clustered in module 149 of the colon-transverse network, enriched for TRAIL-activated apoptotic signaling pathway genes, a long-known signaling pathway involved in cancer progression (42) (Fig. 3C). Breast and ovarian cancer heritability are also clustered in modules involved in estrogen metabolism and signaling (breast cancer and module 180 of skin–sun-exposed, ovarian cancer and module 182 of skin–not sun-exposed).

Metabolic traits such as high blood levels of HDL (high-density lipoprotein) and Type 2 Diabetes tend to cluster in modules enriched for lipid and carbohydrate metabolism. High HDL level heritability is enriched in module 142 of adipose–visceral omentum, enriched for genes involved in very-low and high-density lipoprotein particle assembly, remodeling, and clearance (Fig. 3D). Type 2 diabetes heritability clusters in module 17 of whole blood, enriched in genes involved in glycogen metabolism (Fig. 3F). Finally, heritability for height, a trait primarily related to development and, in particular, bone and muscle growth, is clustered in module 90 of muscle–skeletal, enriched for genes involved in muscle tissue morphogenesis, endochondral ossification, bone mineralization, and chondrocyte and osteoblast differentiation (Fig. 3E).

### Local and global hubs carry a large part of trait heritability

We then explored how heritability is distributed across the SNPs nodes in the eQTL network and whether there are particular network topological features that correlate with a greater-than-expected ability to explain the heritability of a particular trait. We thus computed two different summary statistics characterizing the topological properties of SNPs within each network: outdegree and core score (24). The outdegree measures the centrality of a SNP within the entire network. SNPs within the top 25% of outdegree distribution were considered as global hubs. The core score measures the contribution of SNPs to the modularity of the network module in which it arises. SNPs within the top 25% of core score distribution were considered local hubs (core SNPs).

We investigated whether trait heritability was distributed evenly across all SNPs or instead concentrated in SNPs with local or global centrality. Using a likelihood ratio test accounting for linkage disequilibrium and module size (see Methods), we found for almost every trait we considered that the SNPs explaining the greatest portion of the heritability (top 5% of *Z*^2^, or high heritability SNPs) are more likely to have both high outdegree and core scores (Supplementary Fig. S2 for outdegrees and Supplementary Fig. S3c for core scores). The enrichment in high heritability among local “hubs” appears stronger and more systematic than for global “hubs;” in Artery Coronary and Liver, the global hubs do not show any significant enrichment at all. Using the Whole-Blood network as a model, we found that enrichment of high heritability SNPs in local hubs increases with the threshold chosen to define high outdegree and core score (from the top 90% to the top 5%, Supplementary Figure S4A.).

Given the demonstrated importance of eQTL network topology, we explored whether the increased heritability we found was driven simply by being in the network (a proxy for being an eQTL), being a global hub, or being a local hub. These annotations are overlapping and reflect potentially confounding factors such as underlying chromatin annotations (for example, global hubs are enriched for non-genic enhancers, while local hubs are enriched for genic enhancers and promoters (23)). For this reason, we partitioned heritability across various functional annotations while accounting for linked markers using stratified LD Score regression(43, 44) (see Methods) using the 97-levels baseline annotation model, to which we added our three annotations: belonging to eQTL network, being in the top quartile of outdegrees (global hubs), or of core scores (local hubs). The total proportion of heritability explained by SNPs as estimated using the LDSC software is reported in Supplementary Tab. S2. We found that these proportions are not always perfectly correlated with the estimated global genetic heritability reported in published twin and pedigree studies (see Supplementary Tab. S2), but our estimate of the total heritability of traits explained by SNPs as computed by the LD score regression is coherent with previous reports (see (45) for breast, ovarian and prostate cancer examples).

The LD score regression confirmed the results we obtained above using likelihood ratio tests: SNPs with high core scores or high outdegrees are enriched for trait heritability, and this enrichment increases with the threshold chosen to define high scores (Supplementary Fig. S4B). The detailed results that show enrichment in *h*^2^ among SNPs within each annotation for each of the ten traits and diseases are reported in Supplementary Tab. S8.

We performed a meta-analysis across the ten uncorrelated traits and diseases for each tissue-specific network. An example of enrichment in *h*^2^ among each annotation for the Whole-Blood network is shown in Fig. 4A. As noted earlier, SNPs that are within the eQTL networks tend to be significantly enriched in *h*^2^ independent of tissue (Fig. 4B). However, the enrichment in *h*^2^ is even greater for both high outdegree SNPs and high core score SNPs in all networks except Artery - Coronary and Liver. In many ways, this trend is exactly what one would expect: SNPs that fall within the eQTL networks have increased heritability because they are potentially capable of affecting the expression of multiple genes, a trend that increases as the SNPs become increasingly connected in the eQTL networks.

**Fig. 4.**
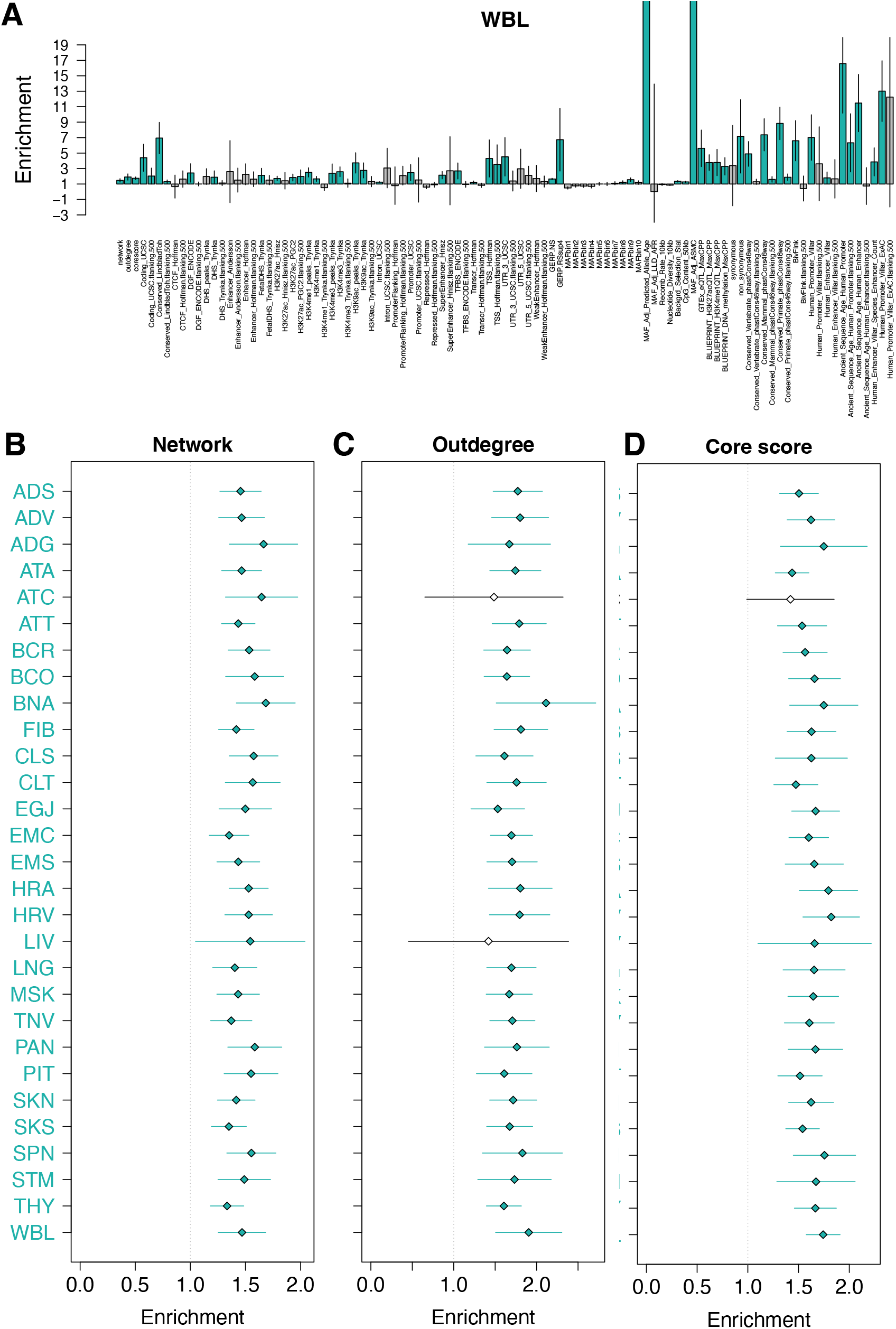
Global and local hubs explain a larger share of heritability than expected by chance. **A**. Meta-analysis of *h*^2^ enrichment among various annotations across 10 uncorrelated traits and diseases for the Whole Blood eQTL network. **B-D**. Meta-analysis of *h*^2^ enrichment among SNP subcategories across 10 uncorrelated traits and diseases for each tissue-specific eQTL network. **B**. Among SNPs within the eQTL network. **C**. Among global hubs (high degrees). **D**. Among local hubs (high core scores.)

### Local hubs evolve under polygenic adaptation

The clustering of complex traits and diseases heritability in a few modules and in hubs in eQTL networks, together with previous results suggesting that global hubs are likely to evolve under negative selection (46) suggest that directional selection on complex traits may preferentially be mediated by local hubs. The importance of the local hubs makes logical sense. Indeed, selection acts at the level of the traits. As each specific trait is likely to be under the control of one or several network modules consisting of genes involved in the same biological function, the local hubs play an important role in their determination. Targeted selection acting on local hubs would moreover mitigate the potential perturbative effects on the network as a whole or other functional modules. To test for this hypothesis, we searched for enrichment among global or local hubs of high scores for different statistics measuring three types of selection signatures. Genomic Evolutionary Rate Profiling score (GERP) detects regions that lack substitutions, which can indicate negative selection (47). Integrated Haplotype Score (*iHS*) detects longer-than-expected haplotypes around one of the alleles at one locus, a signature of recent, strong directional selection event or selective sweeps (48). *F*_*ST*_ measures population differentiation that may reflect older and sometimes milder directional selection events and has proven to be powerful in detecting the shifts in beneficial allele frequencies observed in polygenic selection (49, 50).

We found that both global and local hubs, while enriched for complex traits and disease heritability, were more likely to be located in constrained genomic regions with high GERP scores in all the tissue-specific networks (see Fig. 5A and Supplementary Fig. S5A). Global hubs were even more likely than local hubs to belong to constrained regions S6). Both types of hubs are globally evolving under negative selection, in particular global hubs.

**Fig. 5.**
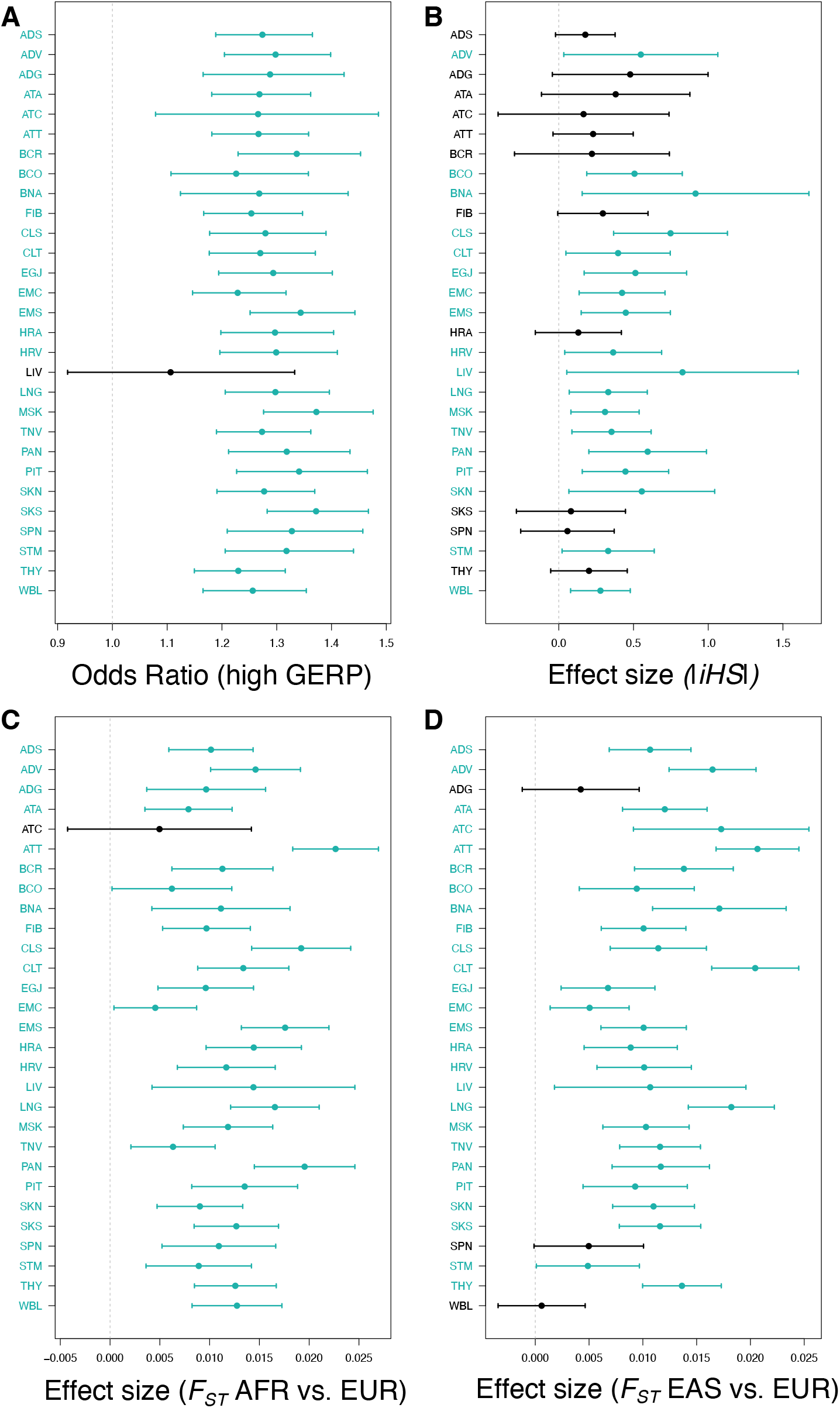
While overall under negative selection, local hubs are enriched for positive selection signals in almost all tissue-specific eQTL networks. **A**. Odds ratio of being conserved when the SNP is a local hub *vs*. a leaf. Bars indicate confidence intervals. Confidence intervals and *p*-values on the left side were computed using a logistic regression. **B-D**. Effect size of being a local hub vs a leaf on the value of several statistics sensitive to positive selection. Bars indicate 2 *× SE*. Standard error (*SE*) and *p*-values on the left side were computed using a linear regression. **B**. *iHS*, that detect recent selective sweeps signals. **C**. *F*_*ST*_ (*AFR, EUR*) measuring population differentiation between African (AFR) and European (EUR) samples from 1000 genomes. **D**. *F*_*ST*_ (*EAS, EUR*) measuring population differentiation between East-Asian (EAS) and European (EUR) samples from 1000 genomes. **A-D**. For each tissue-specific network, significant values (an odds ratio significantly different from 1 or effect size significantly different from 0 are highlighted in blue. Significance was called when *p* ≤ 0.05.

Neither local nor global hubs presented a clear pattern of enrichment for high |*iHS*| in all tissue-specific networks (see Fig. 5B and Supplementary Fig. S5B). However, local hubs were significantly enriched for high |*iHS*| in more tissue-specific networks than global hubs (18 of the 29 networks for local hubs *vs*. 13 of the 29 networks for global hubs, see Fig. 5B and Supplementary Fig. S5B). Local hubs were also enriched for high *F*_*ST*_ between European and both African and East Asian populations in almost all networks. Conversely, global hubs were depleted for high *F*_*ST*_ in almost all networks. Local hubs thus showed more signatures corresponding to polygenic selection than the other SNPs in the network, while global hubs seemed to be evolving under strict negative selection.

## Discussion

Of the SNPs found through GWAS to be associated with complex traits, most (up to 90%) fall outside of gene coding regions suggesting that they likely play a regulatory role. Bipartite eQTL networks have allowed us to explore how these SNP loci work together to regulate the many genes involved in the biological processes underlying the traits, in a tissue- or cell-type-specific manner (23, 24, 26). Using eQTL networks, we found that not only do SNPs act in both *cis-* and *trans-* to influence the expression of complex networks of genes, but that these networks have a robust, modular structure consisting of SNP-gene modules with properties that help explain the polygenic determinism that underlie most common traits and provide clues on how they can evolve despite the high degree of pleiotropy. Specifically, the modules in eQTL networks are enriched for functionally related groups of genes and nucleated around “core SNPs” that are both local (module) hubs and the SNPs most likely to exhibit strong GWAS associations with various phenotypes (including diseases) due to their association with module-driven, trait-related processes (23, 24).

The omnigenic model was important in that it helped bridge the gap between the missing heritability found in many diseases and the growing number of traits for which hundreds, if not thousands, of small effect-size genetic variants contribute to a given trait. Our eQTL network-based model is complimentary to the omnigenic model but provides a more nuanced view of the importance of SNP-gene “regulatory” associations in defining phenotypes, identifying disease associations and functions, and explaining both missing heritability and selection. Although it has been reported that global hubs in eQTL networks are enriched for tissue-relevant trait heritability (25), as are gene modules appearing in various types of networks(22), there has not been a systematic exploration of these two complementary and important conceptual advances in understanding genetic effects in complex traits.

We addressed that gap in understanding by performing an analysis of ten polygenic traits that represent a wide range of genetic heritability to determine the distribution of trait heritability across 29 tissue-specific eQTL networks. We found that although heritability is widely distributed across loci, as suggested by the omnigenic model, the distribution of heritability is far from homogeneous. Instead, there is an uneven distribution with the greatest heritability concentrated in network modules containing genes that represent trait-specific and biologically relevant functions. This makes sense as phenotypic traits arise through alterations of specific biological processes relevant to individual traits. Further, we found that trait heritability was more likely to be explained by SNPs occupying key positions in the eQTL networks, especially among the “core SNPs” that are local hubs in their functional modules. These same SNPs, which we had previously shown to be enriched in tissue-specific activated regulatory elements(23), are thus likely to determine a significant proportion of the heritability of complex traits.

The clustering of heritability in modules is also not evenly distributed across traits and differs among tissues. We found that heritability in each phenotype tended to be clustered in a small number of biologically relevant modules within the highly modular eQTL networks that are relevant for understanding the phenotype in question. As previously noted, these modules tend to be trait-specific, even when traits are not genetically correlated. This module-dependent concentration of heritability, coupled with the enrichment of heritability in core SNPs of those same modules, could help explain why even highly connected regulatory networks (19, 20) are robust to the genetic perturbations of deleterious genetic variants. Indeed, the highly modular structure of eQTL networks provides a means by which the disruptive effect of regulatory mutations can be buffered in a tissue-specific manner against altering the broader functionality of the wider regulatory networks active in living cells.

We also found that global hubs are evolving under strong constraints, as observed in previous studies (46). However, our results revealed that another category of nodes, the local hubs that regulate many genes involved in the same biological processes and articulate modules, have been preferential targets of past and ongoing polygenic selection. Because of their strategic position in the eQTL regulatory network, these loci greatly influence the regulation of biological processes in a tissue-specific manner, and, because their effect is limited beyond their “home” module, they are ideal candidates for adaptation. Finally, our results confirm the hypothesis, first presented in an opinion paper by Fagny and Austerlitz (51), that local hubs are preferential targets for polygenic selection, and provide a means to understand how complex traits may evolve despite the strong pleiotropy that exists in biological networks.

Overall, our results demonstrate a synergy between the eQTL network and omnigenic models in explaining how genetic variants work together to influence traits while addressing their respective shortcomings. The value of such a synthesis can be seen in the results derived from the collection of complex traits we chose to analyze, including cancers, metabolic diseases, and auto-immune neurodegenerative disorders. In each of these traits we can see genetic risk factors identified through GWAS perturbing tissue-specific functional modules (while affecting others), while those modules are simultaneously affected by many other genetic variants of smaller overall effect size. These complex relationships define a conceptual framework for understanding disease risk, helps to define phenotypes for health and disease, and provide a means by which one could potentially prioritize therapeutic targets and design treatment protocols that account for the network architecture. Our findings also allow a better understanding of how complex traits can change as populations adapt to local changes through selection acting on traits mediated at the genetic and molecular level by key regulatory elements. This conceptual framework also provides an explanation, at the molecular level, of how polygenic selection of regulatory mutations beneficial in one environment could lead to the phenomenon of maladaptation that many suspect is the origin of many complex diseases. Finally, if the lessons we learned here were transferable to other organisms, they could be of tremendous value to breeders by pointing out which loci are the more likely to improve the tolerance of crops or cattle to environmental stresses without jeopardizing their agricultural values.

## Author Contributions

KS, JP, JQ, and MF contributed to the conception of the work. KS, JP, JQ, and MF contributed to the study design and method development. KS, JP, and MF contributed analysis, verified the data, and drafted the manuscript. All authors reviewed the manuscript and approved the submitted work.

## Ethics approval

This work was conducted under dbGaP-approved protocol #9112.

## ACKNOWLEDGEMENTS

Thank you to Alkes Price for discussions about LDSC and heritability and for making the GWAS summary statistics data available. The Genotype-Tissue Expression (GTEx) Project was supported by the Common Fund of the Office of the Director of the National Institutes of Health, and by NCI, NHGRI, NHLBI, NIDA, NIMH, and NINDS. This work was supported by grants from the US National Institutes of Health, including grants from the National Heart, Lung and Blood Institute (5P01HL105339, 5P01HL114501; J.Q. and J.P.: 5R01HL111759; J.P.: K25HL140186), the National Cancer Institute (J.Q.: R35CA220523, 5P30CA006516; J.Q. and M.F.: 1R35CA197449), the National Institute of Allergy and Infectious Disease (J.Q. and J.P.: 5R01AI099204), the National Human Genome Research Institute (J.Q.: R01HG011393); and the Marie Sklodowska-Curie grant PATTERNS (M.F.: 845083).

## Supplementary Information

### Supplementary Text

#### eQTL networks are highly structured

Using RNA-seq and genotyping data from 29 tissues with *>* 200 matching samples from the GTEx data, we performed eQTL studies (see Methods). Including all associations with a Benjamini-Hochberg corrected *p*-value *<* 0.2, we built 29 tissue-specific weighted bipartite eQTL networks, using 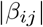, the coefficient of the linear regression for SNP *i* and gene *j* as weights.

The structure of each eQTL network in terms of modules was then inferred using a new implementation of the bipartite maximization algorithm that balances computation time and memory usage (see Methods). All of the networks are highly structured, with modularity values ranging from 0.92 to 0.98 (Supplementary Fig. S1A). To note, as previously shown (23), the eQTL network structure does not recapitulate local linkage disequilibrium, but rather summarizes the genetic component of gene expression regulation across the genome, with most modules presenting genes and SNPs from at least 2 different chromosomes (Supplementary Fig. S1B). The description in terms of the number of nodes (SNPs and genes) and edges can be found in Supplementary Fig. S1C-E.

### Supplementary Tables

**Table S1.**
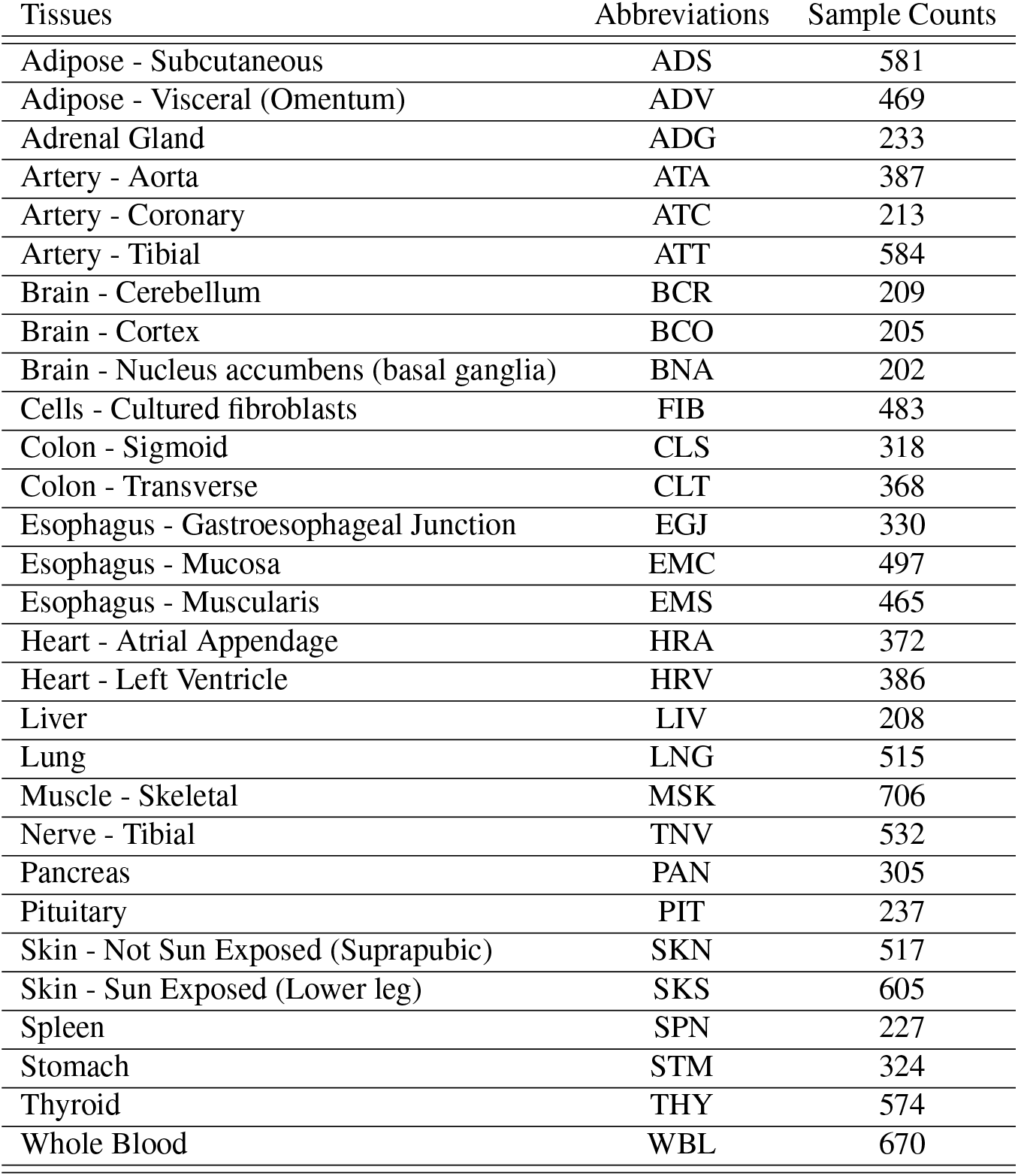
Description of GTEx tissues.

| Tissues | Abbreviations | Sample Counts |
| --- | --- | --- |
| Adipose - Subcutaneous | ADS | 581 |
| Adipose - Visceral (Omentum) | ADV | 469 |
| Adrenal Gland | ADG | 233 |
| Artery - Aorta | ATA | 387 |
| Artery - Coronary | ATC | 213 |
| Artery - Tibial | ATT | 584 |
| Brain - Cerebellum | BCR | 209 |
| Brain - Cortex | BCO | 205 |
| Brain - Nucleus accumbens (basal ganglia) | BNA | 202 |
| Cells - Cultured fibroblasts | FIB | 483 |
| Colon - Sigmoid | CLS | 318 |
| Colon - Transverse | CLT | 368 |
| Esophagus - Gastroesophageal Junction | EGJ | 330 |
| Esophagus - Mucosa | EMC | 497 |
| Esophagus - Muscularis | EMS | 465 |
| Heart - Atrial Appendage | HRA | 372 |
| Heart - Left Ventricle | HRV | 386 |
| Liver | LIV | 208 |
| Lung | LNG | 515 |
| Muscle - Skeletal | MSK | 706 |
| Nerve - Tibial | TNV | 532 |
| Pancreas | PAN | 305 |
| Pituitary | PIT | 237 |
| Skin - Not Sun Exposed (Suprapubic) | SKN | 517 |
| Skin - Sun Exposed (Lower leg) | SKS | 605 |
| Spleen | SPN | 227 |
| Stomach | STM | 324 |
| Thyroid | THY | 574 |
| Whole Blood | WBL | 670 |

**Table S2. Description of GWAS data**. See GWASDescription.txt.

**Table S3. Enrichment of each module in high mean** *Z*^2^ **and high heritability SNPs**. See all_signif_enrichment_bycluster.txt.

**Table S4. Number of modules enriched with per-SNP** |*Z*^2^ **and high heritability SNPs for each trait / tissue-specific network pair**. See summary_signif_enrichment_bycluster.txt.

**Table S5. Pairwise comparison between traits of modules enriched in high heritability SNPs**. See overlapping_enrichment_bycluster_FDR.txt.

**Table S6. Gene Ontology enrichment analysis results for each module in each tissue-specific network**. See topGO_results.txt.

**Table S7. Gene Ontology enrichment analysis results for each module in each tissue-specific network**. See tissue-specific_trait-specific_enriched_modules.txt.

**Table S8. Enrichment of** *h*^2^ **in various annotations obtained with LDSC** See results_LDSCORE_all_scores.txt.

### Supplementary Figures

**Fig. S1.**
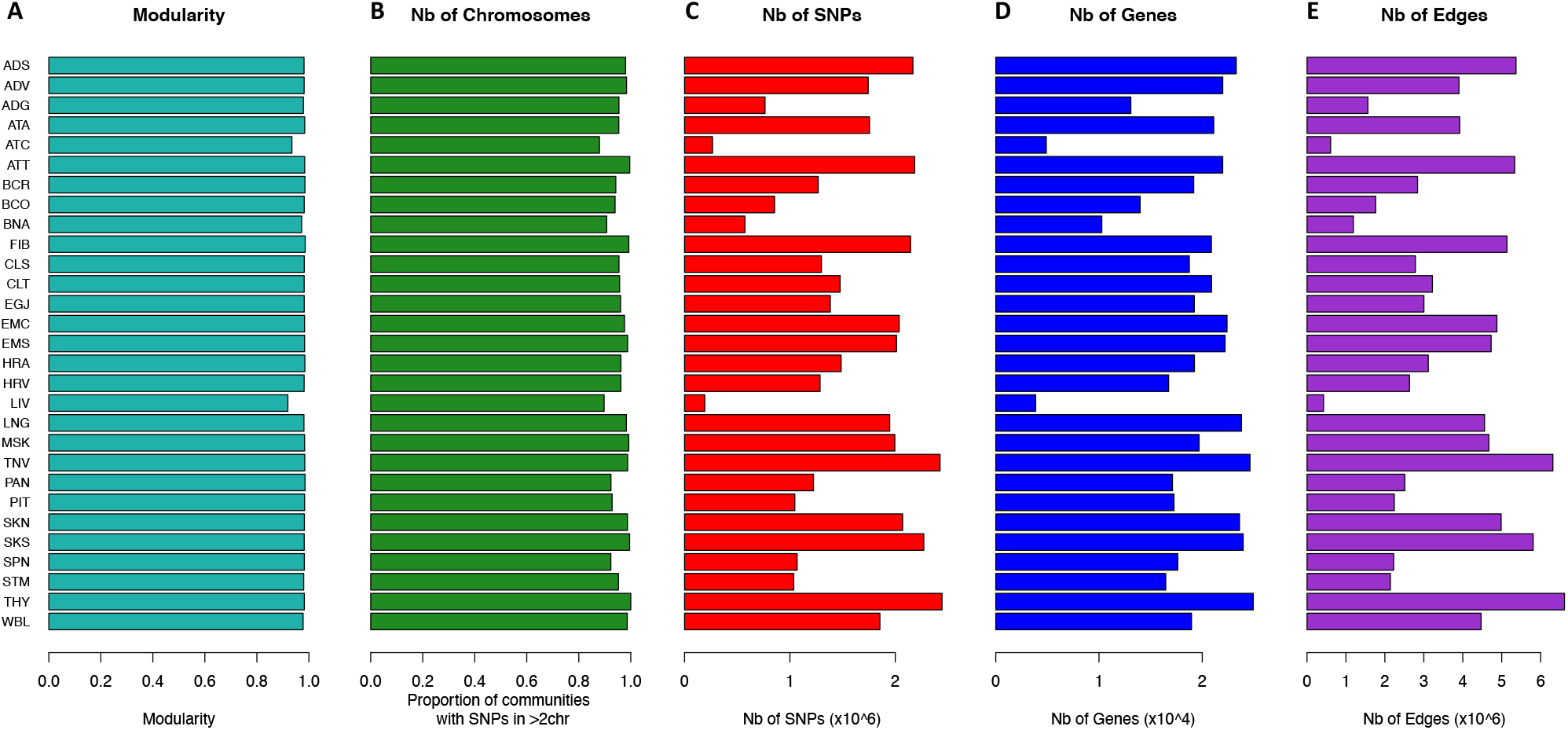
eQTL networks are highly structured and reflect regulatory relationships. **A**. Modularity of each tissue-specific eQTL network. **B**. Most modules from eQTL networks contain SNPs and genes from several modules and are not driven by linkage disequilibrium. **C-E**. Number of nodes and edges of each tissue-specific eQTL network. **C**. SNPs. **D**. Genes. **E**. Edges at FDR < 0.2.

**Fig. S2.**
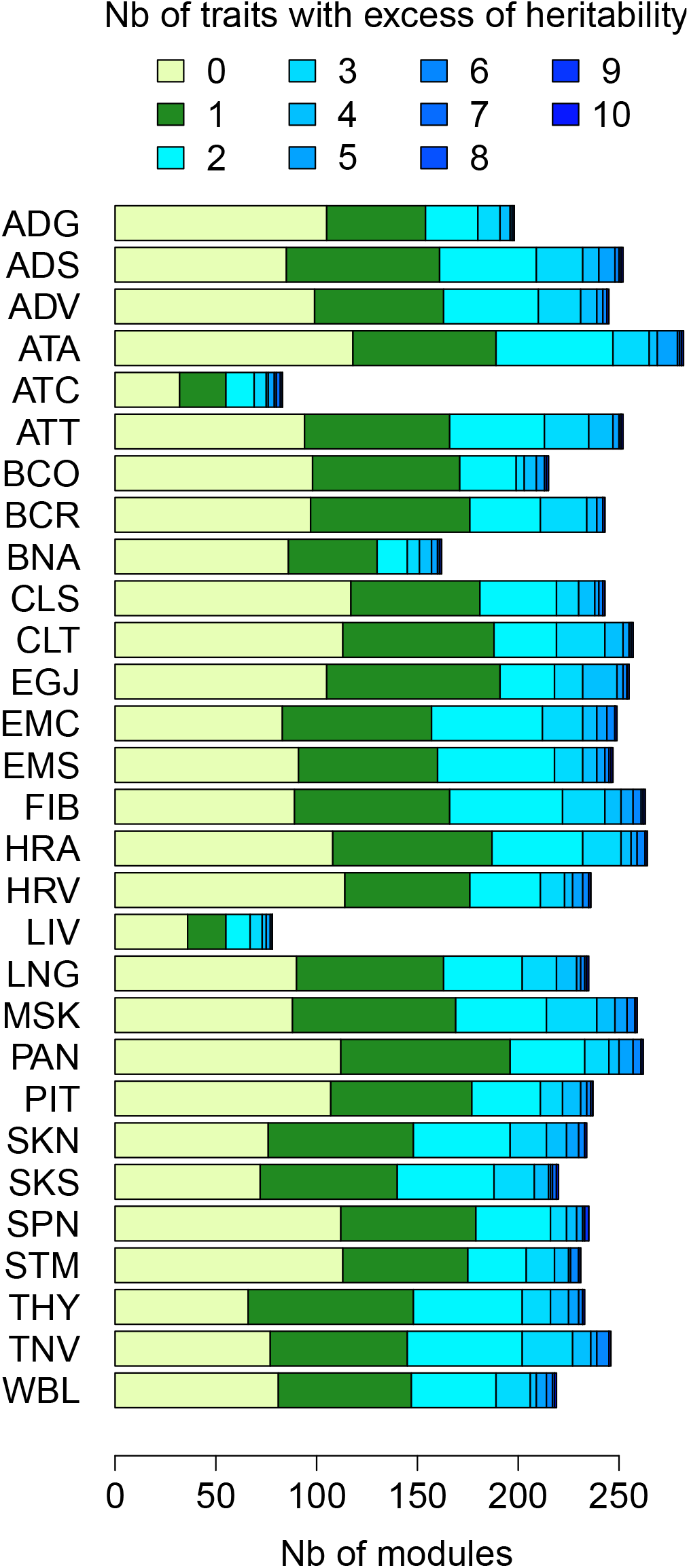
Proportion of modules enriched for heritability for 0 to 10 traits.

**Fig. S2.**
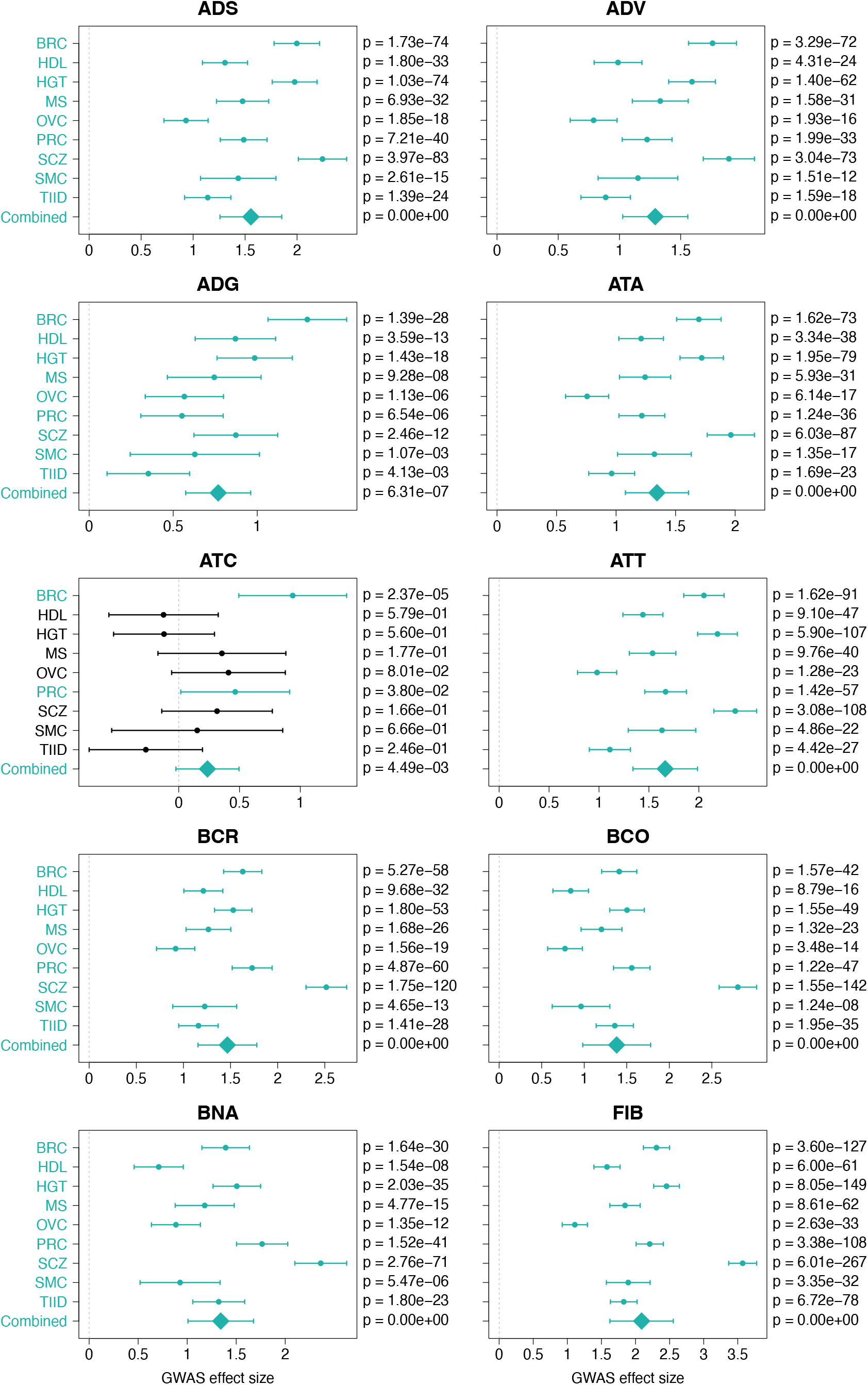
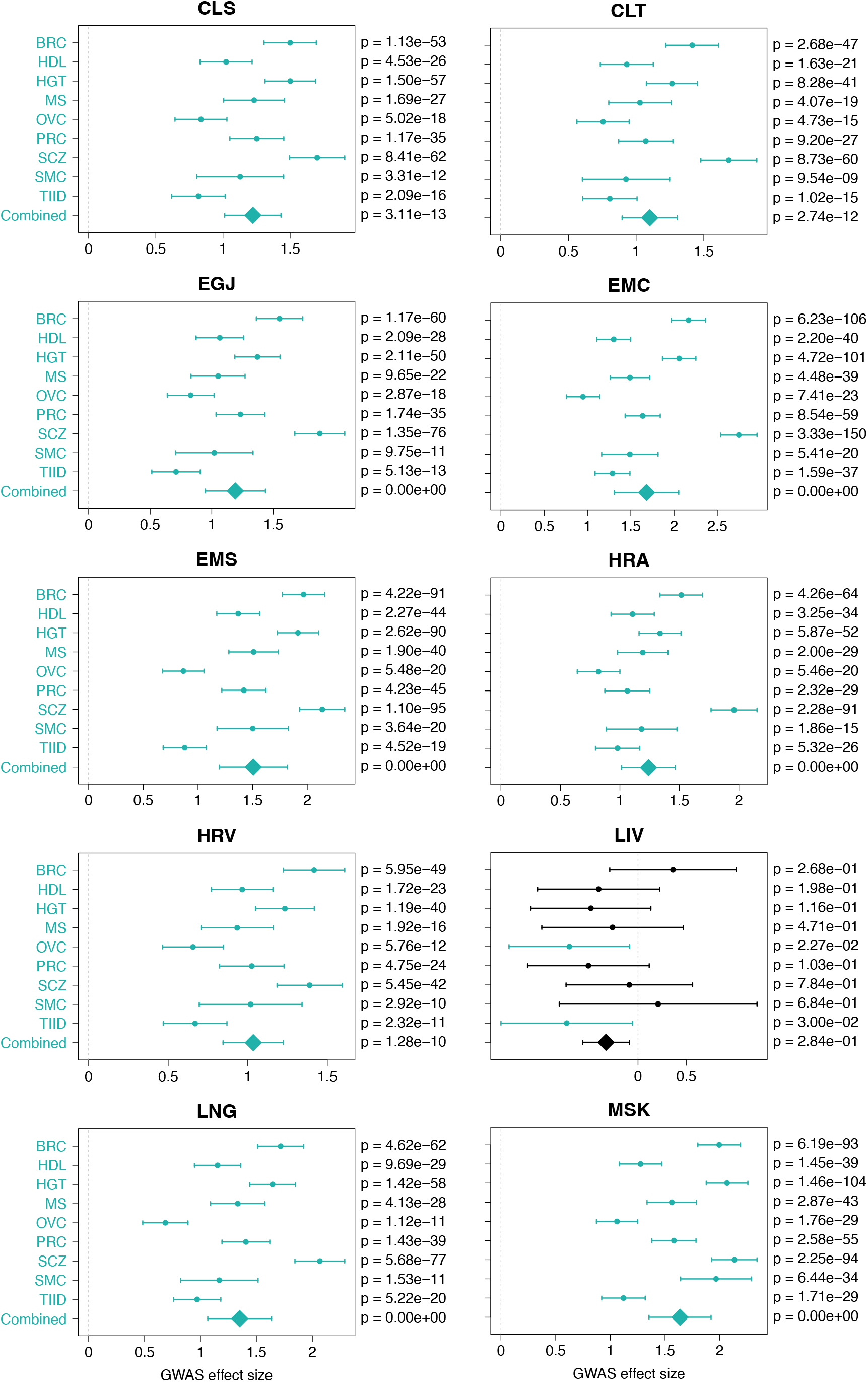
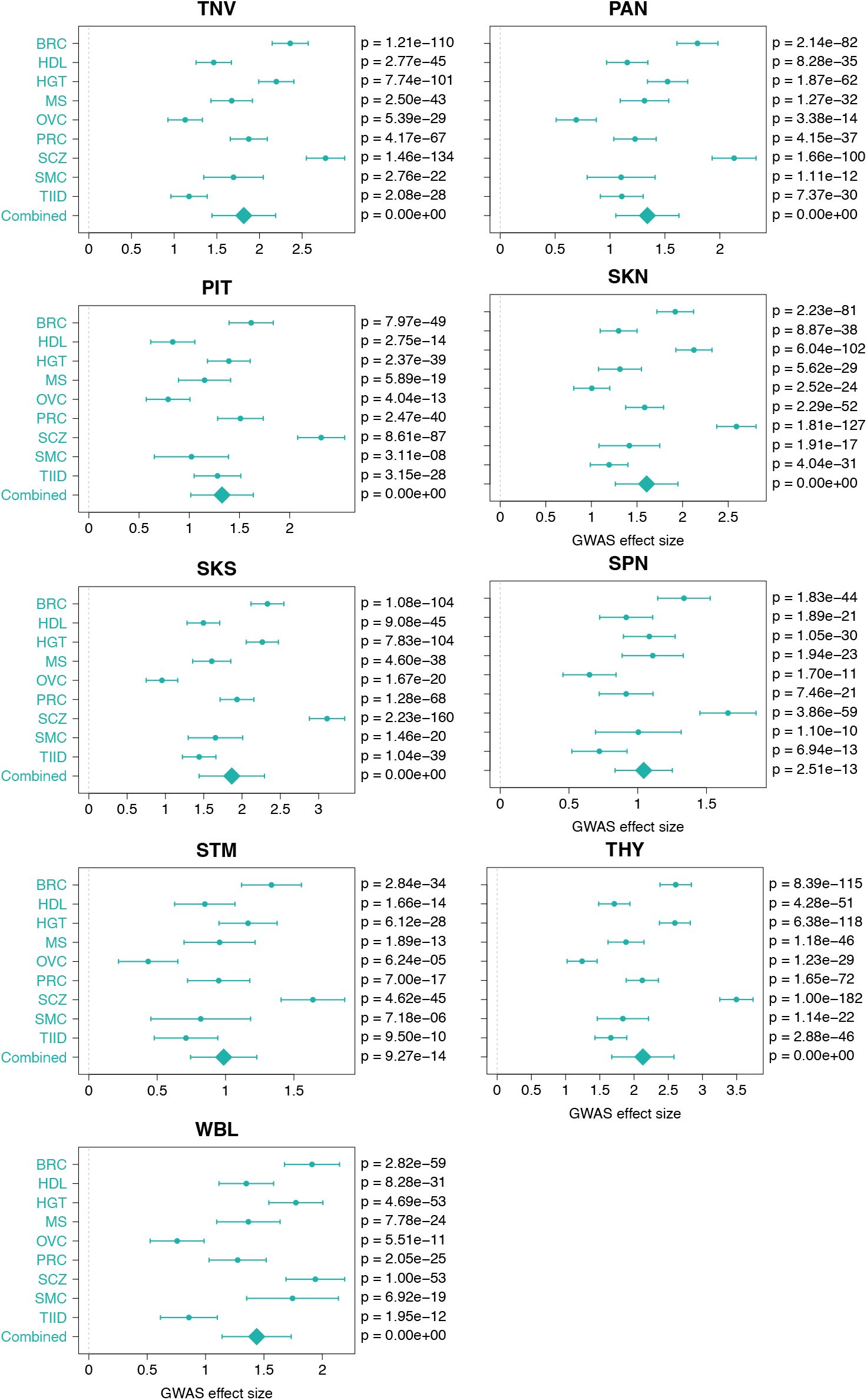
High heritability SNPs have network higher degree. Effect size of being a high heritability SNP on outdegree in each of the tissue-specific networks. *p*-values were obtained using a likelihood ratio test correcting for linkage disequilibrium.

**Fig. S3.**
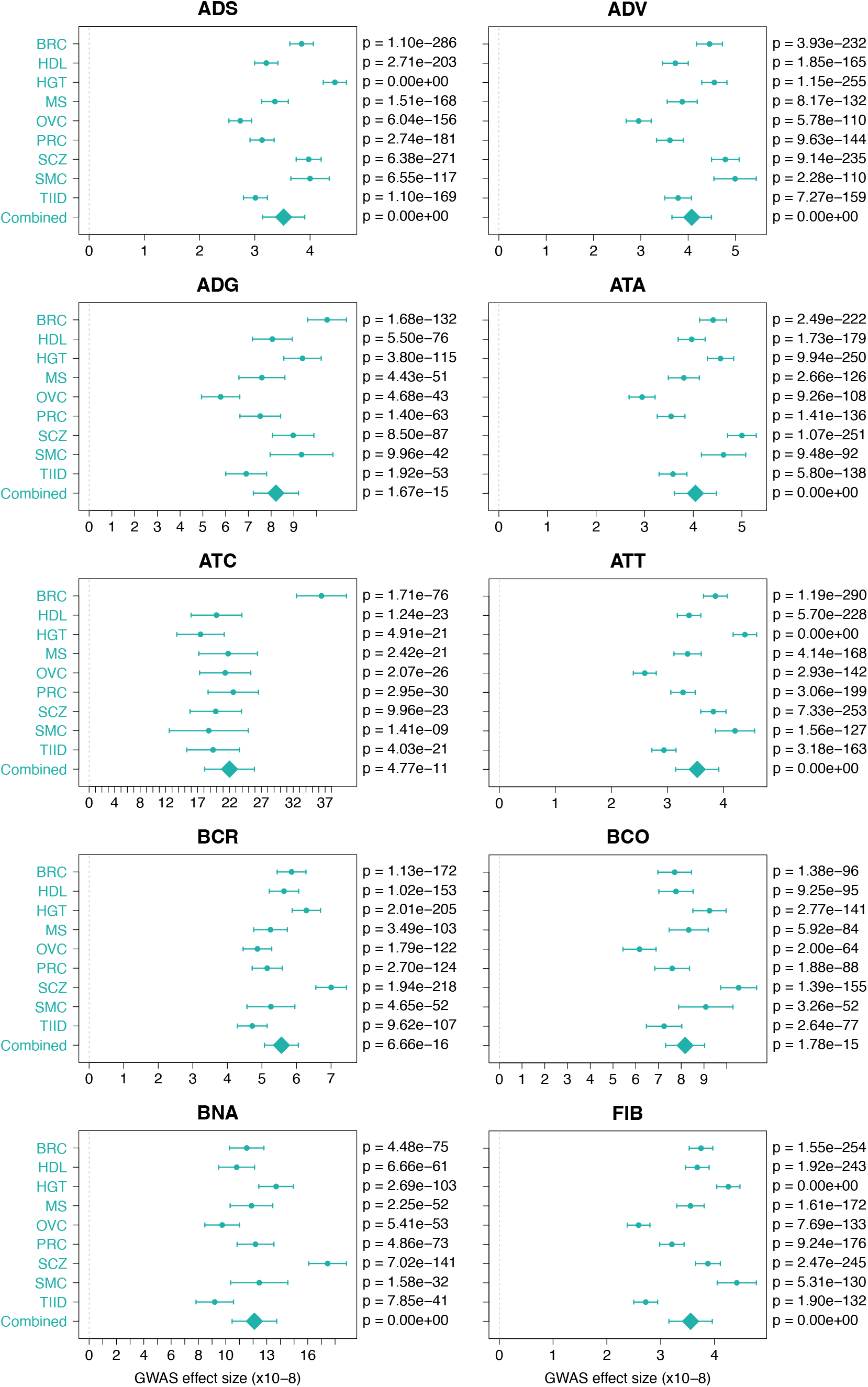
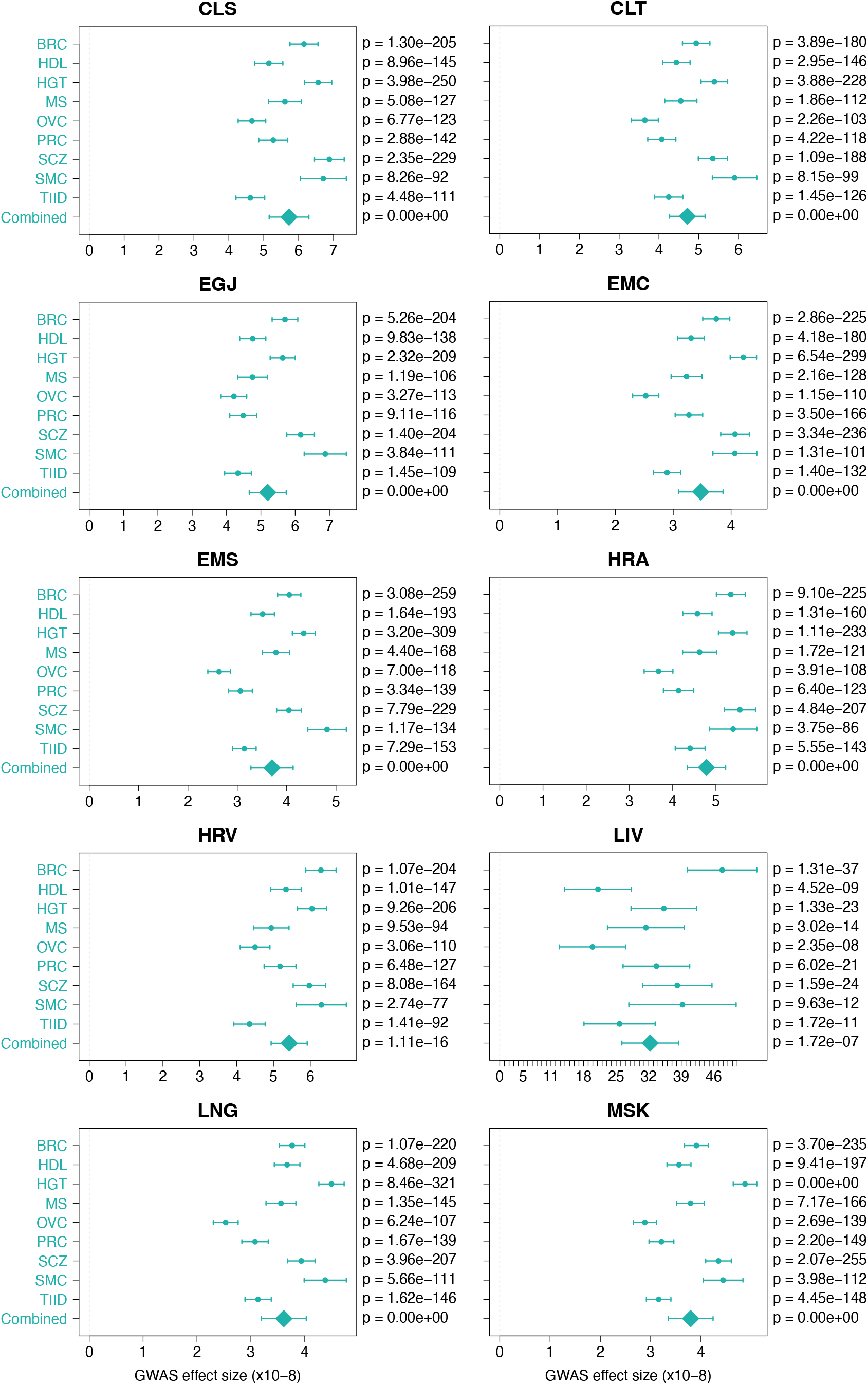
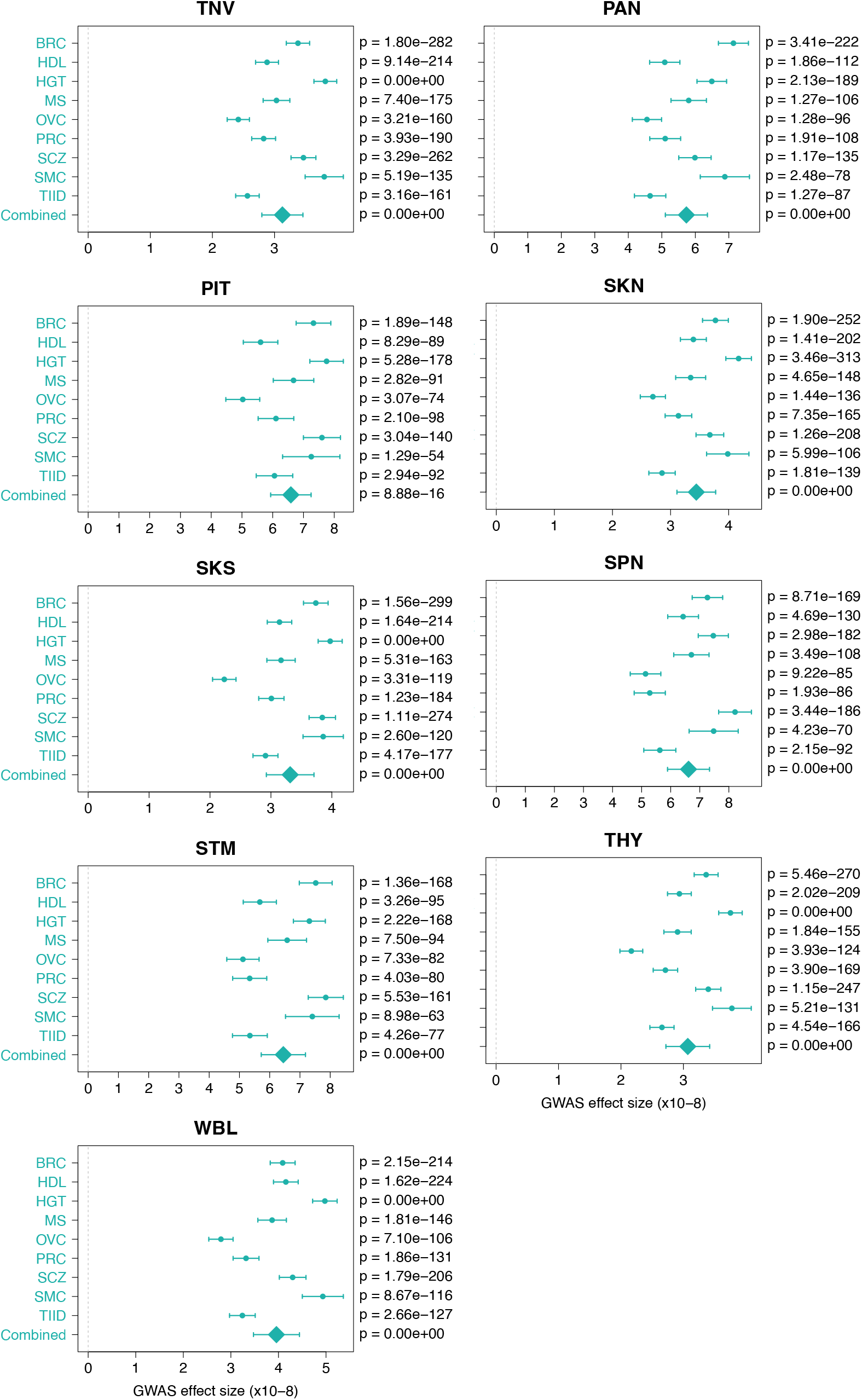
High heritability SNPs have higher core scores. Effect size of being a high heritability SNP on core score in each of the tissue-specific networks. *p*-values were obtained using a likelihood ratio test correcting for linkage disequilibrium and module size. As core scores depend on the number of nodes in a module, core scores were corrected for module identity.

**Fig. S4.**
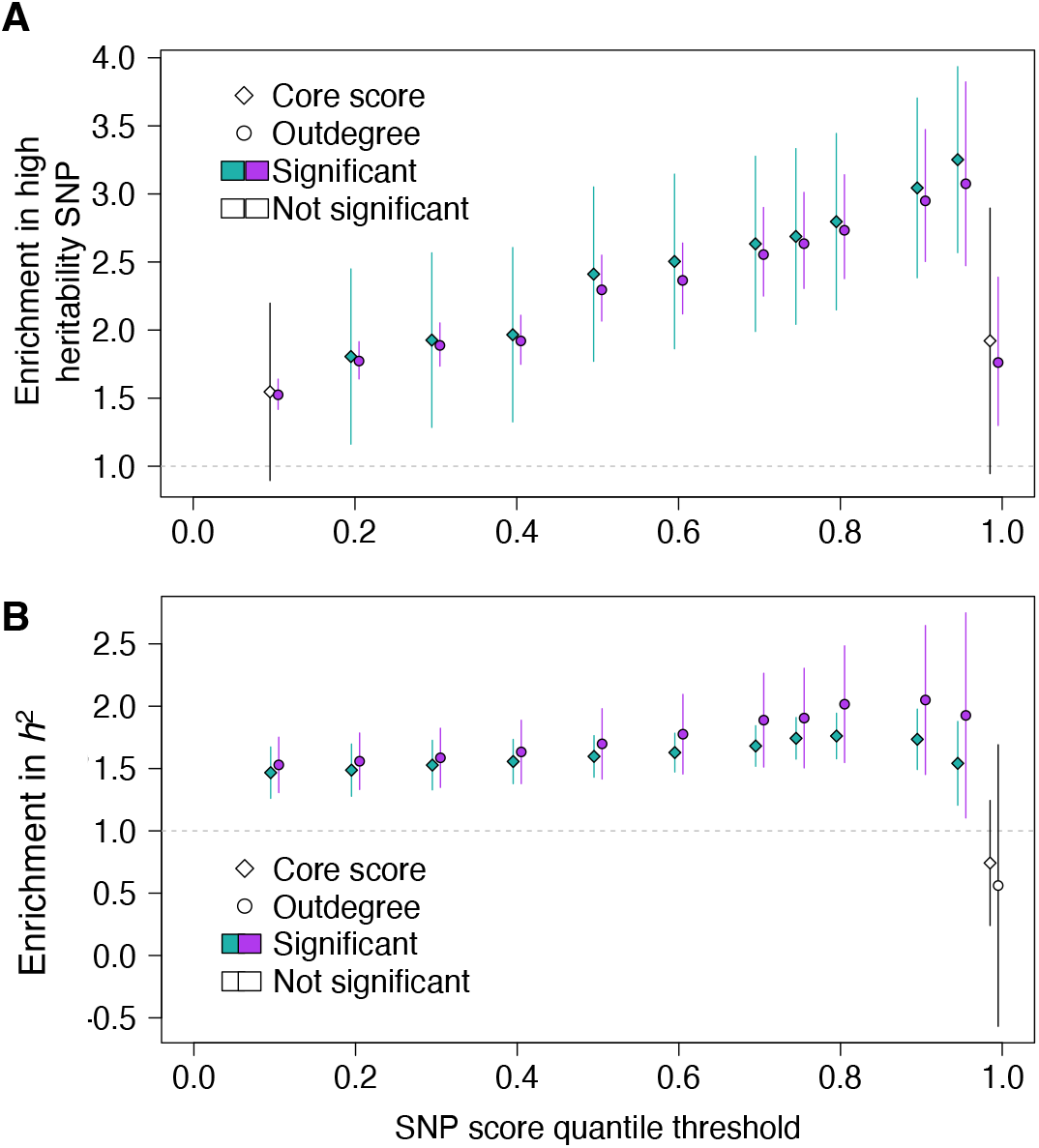
Local and global hubs explain a larger share of heritability than expected by chance. Enrichments were computed using different thresholds to define high core scores and high outdegree in the whole-blood eQTL network from the first 9 deciles to the top percentile. **A**. Enrichment of high heritability SNPs among local and global hubs obtained using a likelihood ratio test. **B**. Enrichment of *h*^2^ explained by global and local hubs obtained using stratified LD Scores and the basal model. **A-B**. Diamonds = core scores (local hubs). Circles = degrees (global hubs). Colored points = significant enrichment computed using a meta-analysis across 10 uncorrelated traits and diseases.

**Fig. S5.**
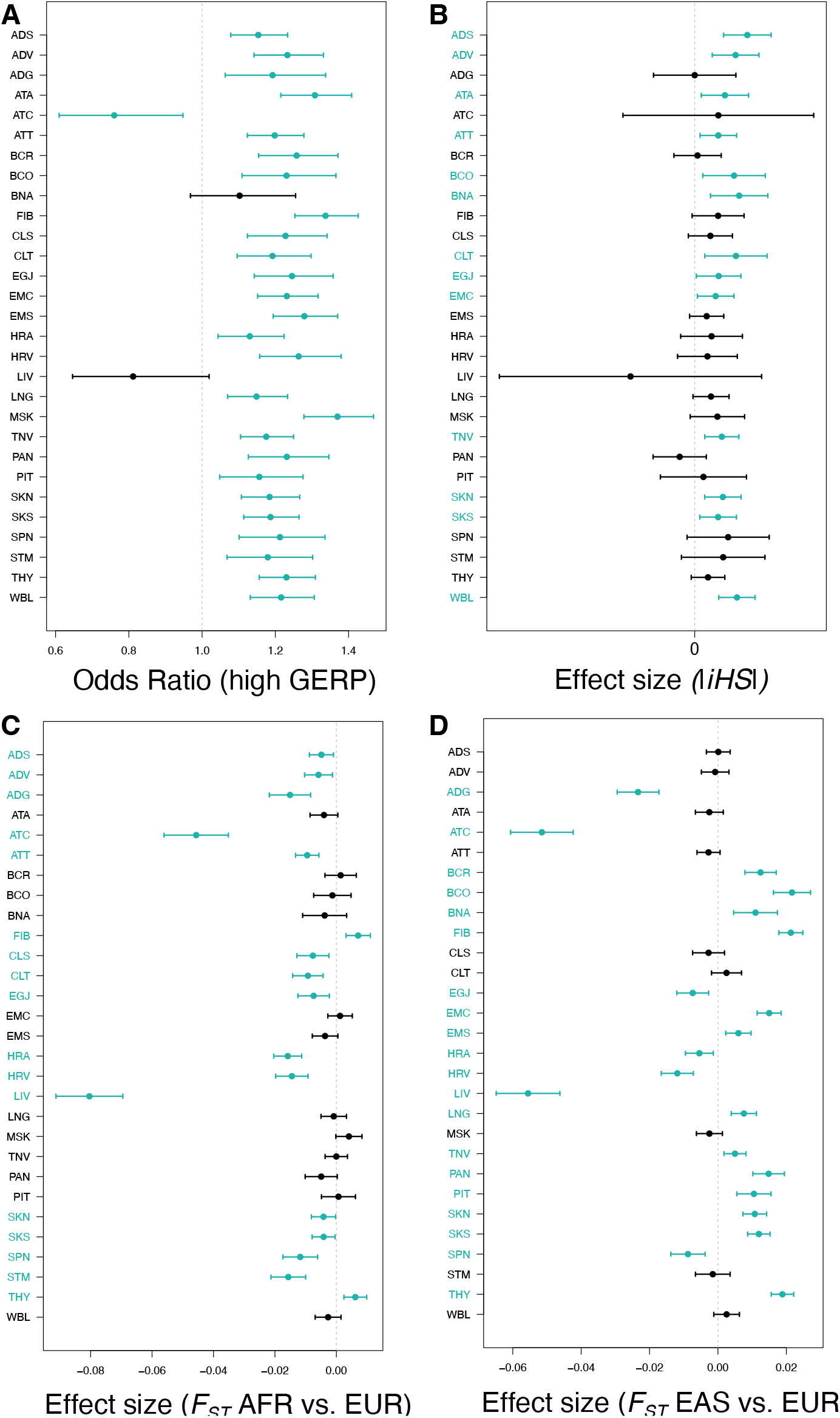
Global hubs are mainly evolving under negative selection. **A**. Odds ratio of being conserved when the SNP is a local hub *vs*. a leaf. Bars indicate confidence intervals. Confidence intervals and *p*-values on the left side were computed using a logistic regression. **B-D**. Effect size of being a local hub vs a leaf on the value of several statistics sensitive to positive selection. Bars indicate 2 *× SE*. The standard error (*SE*) and *p*-values on the left side were computed using linear regression. **B**. *iHS*, that detect recent selective sweeps signals. **C**. *F*_*ST*_ (*AFR, EUR*) measuring population differentiation between African (AFR) and European (EUR) samples from 1000 genomes. **D**. *F*_*ST*_ (*EAS, EUR*) measuring population differentiation between East-Asian (EAS) and European (EUR) samples from 1000 genomes. **A-D**. For each tissue-specific network, significant values (an odds ratio significantly different from 1 or effect size significantly different from 0) are highlighted in blue. Significance was called when *p* ≤ 0.05.

**Fig. S6.**
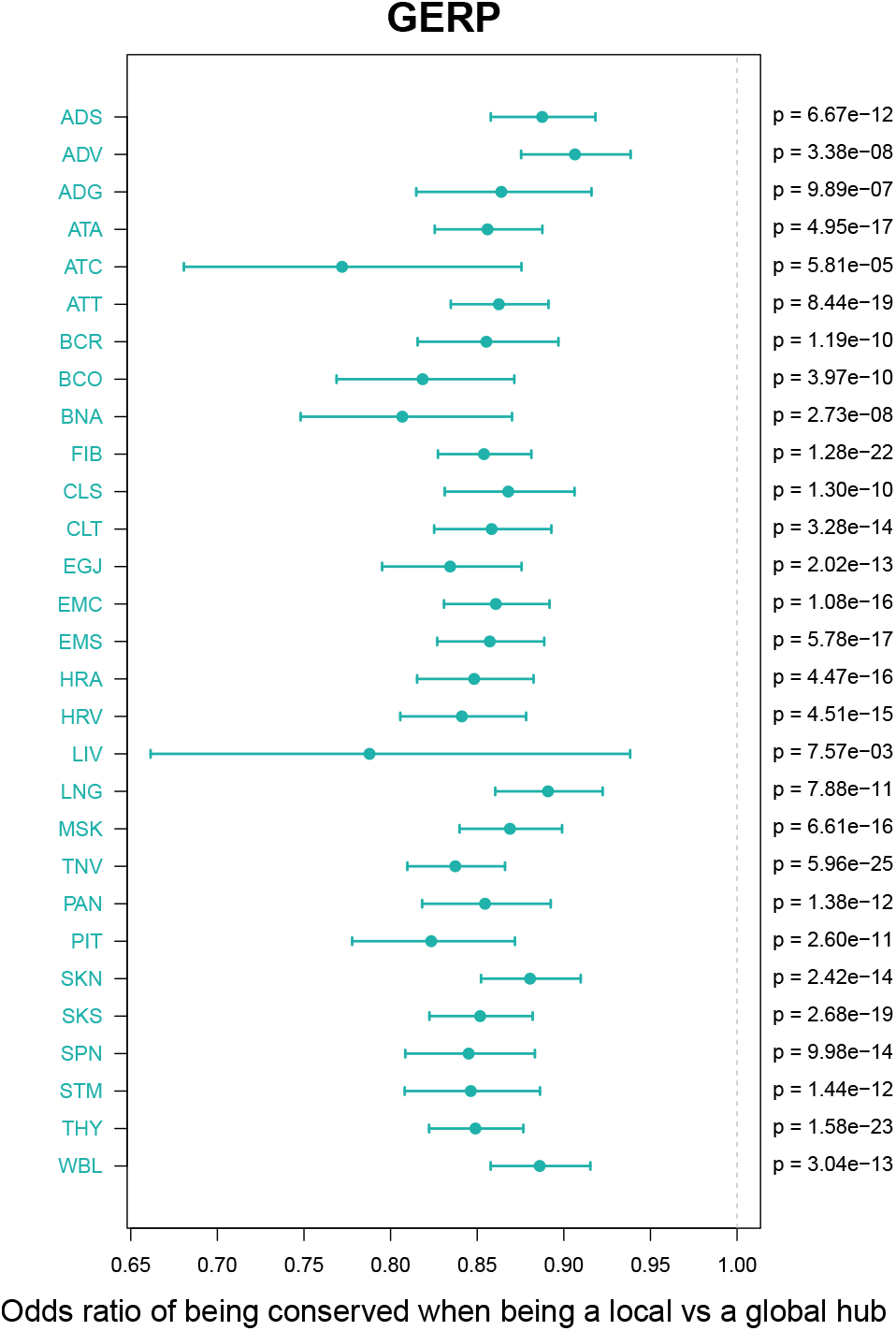
Local hubs are less likely to be under negative selection than global hubs. Odds ratio of being conserved when the SNP is a local hub *vs*. a global hub in each tissue-specific eQTL network. Bars indicate confidence intervals. Confidence intervals and *p*-values on the left side were computed using a logistic regression. Significant values (an odds ratio significantly different from 1) are highlighted in blue. Significance was called when *p* ≤ 0.05.

